# Scrutinizing the SARS-CoV-2 protein information for the designing an effective vaccine encompassing both the T-cell and B-cell epitopes

**DOI:** 10.1101/2020.03.26.009209

**Authors:** Neha Jain, Uma Shankar, Prativa Majee, Amit Kumar

## Abstract

Novel SARS coronavirus (SARS-CoV-2) has caused a pandemic condition world-wide and has been declared as public health emergency of International concern by WHO in a very short span of time. The community transmission of this highly infectious virus has severely affected various parts of China, Italy, Spain and USA among others. The prophylactic solution against SARS-CoV-2 infection is challenging due to the high mutation rate of its RNA genome. Herein, we exploited a next generation vaccinology approach to construct a multi-epitope vaccine candidate against SARS-CoV-2 with high antigenicity, safety and efficacy to combat this deadly infectious agent. The whole proteome was scrutinized for the screening of highly conserved, antigenic, non-allergen and non-toxic epitopes having high population coverage that can elicit both humoral and cellular mediated immune response against COVID-19 infection. These epitopes along with four different adjuvants were utilized to construct a multi-epitope vaccine candidate that can generate strong immunological memory response having high efficacy in humans. Various physiochemical analyses revealed the formation of a stable vaccine product having a high propensity to form a protective solution against the detrimental SARS-CoV-2 strain with high efficacy. The vaccine candidate interacted with immunological receptor TLR3 with high affinity depicting the generation of innate immunity. Further, the codon optimization and in silico expression show the plausibility of the high expression and easy purification of the vaccine product. Thus, this present study provides an initial platform of the rapid generation of an efficacious protective vaccine for combating COVID-19.

## INTRODUCTION

The most disastrous outbreak of recent days which have put the world into a pandemic threat involves the novel Severe Acute Respiratory Syndrome Coronavirus 2 or the SARS-CoV-2 or the 2019-nCoV causing the COVID-19 outbreak. According to the situation report published by World Health Organization on March 25th, 2020, 413,467 confirmed cases have been reported globally along with 18,433 deaths affecting over 197 countries world-wide (https://www.who.int/emergencies/diseases/novel-coronavirus-2019). Due to easy and efficient transportation and migration throughout the world, the virus has spread to 197 countries or areas from its epicenter, i.e. the city of Wuhan, China[1]. The SARS-CoV-2 belongs to the Coronaviridae family being a close relative to the earlier reported Severe Acute Respiratory Syndrome (SARS) virus and Middle East Respiratory Syndrome (MERS) virus. This virus majorly affects the lower respiratory tract causing pneumonia, fever, dry cough, sore throat, dyspnea like symptoms which resolve spontaneously but certain cases result in fatal complications like severe pneumonia, acute respiratory distress syndrome (ARDS), organ failure, septic shock or pulmonary edema often leading to death[2]. The present situation is getting out of control as we don’t have any drug or vaccine to treat this recently emerged virus and supportive treatment and preventive measures are the only ways to restrict COVID-19outcome[3].

Surfacing of SARS-CoV-2 in China was reported recently on December, 2019 and therefore very limited and naïve information is available regarding its genomic constituency and pathogenesis. Hence, the researchers rely on its close cousin viruses, SARS-CoV and MERS-CoV for better understanding of its mode of infection. Phylogenetic analysis have shown that bat-CoV-RaTG13 displays 96.3% sequence similarity while bat-SL-CoVZC45 and bat-SL-CoVZXC21 exhibit about 88% identity with the SARS-CoV-2 virus but the other two pathogenic coronaviruses, SARS-CoV and MERS-CoV have less sequence identity being merely 79% and 50% respectively[4, 5]. Intriguingly, the basic genomic structure for all these coronaviruses remains the same. The single-stranded positive sense RNA genome of SARS-CoV-2 basically encodes 9860 amino acids translating into several non-structural proteins like replicase (orf1a/b), nsp2, nsp3 and accessory proteins (orf3a,orf7a/b) along with structural proteins including Spike (S), Envelope (E), Membrane (M), and Nucleocapsid (N) proteins[4]. The Spike glycoprotein aids in receptor binding and thereby, helps in the entry of the virus in the host cells, while the Envelope protein and the Membrane protein assist in the virus assembly and packaging of the new virions. On the other hand, the Nucleocapsid protein is essential for the genomic RNA synthesis required for the viral survival and proliferation. The non-structural viral proteins including viral enzymes and other accessory proteins help in the SARS-CoV-2 genome replication and infection[6].

The lessons learnt from the SARS-CoV and MERS-CoV outbreak have helped in better understanding of the situation of the new coronavirus epidemic and pushed researchers to develop vaccine or therapeutic solution to succumb the current situation. Previously several animal models including mouse, Syrian hamster, ferrets as well as non-human primate models like rhesus macaque, common marmoset have been used to study SARS-CoV and MERS-CoV and also for the purpose of vaccine development[7–9]. An effective vaccine should robustly activate both the humoral and the cell-mediated immunity to establish strong protection against the pathogen[10]. The antibody generation by B-cell activation as well as acute viral clearance by T-cells along with virus-specific memory generation by CD8+ T-cells are equally important to develop immunity against the coronavirus[11]. In case of respiratory virus like coronaviruses, the mucosal immunity plays an essential role and therefore route of administration of the vaccine is also important in the context of vaccine development. Vaccines designed against previously reported SARS-CoV and MERS-CoV majorly focuses on the Spike protein of virus which consists of S1 and S2 subunits bearing the receptor binding domain (RBD) of the virus and cell fusion machinery respectively. Mutations in the spike protein have also been reported to be responsible for the change in host cell tropism[12]. The S protein is considered most antigenic and thereby can evoke immune responses and generate neutralizing antibodies that can block the virus attachment to the host cells[13]. Other viral proteins which were explored for vaccine development include N protein, E protein and the NSP16 proteins[14–18]. Almost all the platforms for vaccine development for SARS-CoV and MERS-CoV have been investigated including the life-attenuated ones, recombinant viruses, sub-unit protein vaccines, DNA vaccines, Viral vector-based vaccines, nanoparticle-based vaccines, etc. which may form the base for the vaccine designing against the newly emerged SARS-CoV-2[10, 19]. Here we have designed a multi-epitope-based vaccine for the SARS-CoV-2 using next generation vaccinology approach where the recently available genome and proteome of the SARS-CoV-2 were maneuvered and a potential vaccine candidate was conceived. Similar strategy was employed previously for SARS-CoV and MERS-CoV[20–25] as well as certain findings are reported for the newly emerged SARS-CoV-2[26–28]. While immunoinformatics techniques was utilized by groups to predict the B-cell and cytotoxic T-cell epitopes in the SARS-CoV-2 surface glycoprotein, N protein[27–29], others have utilized the information to design epitope-based vaccine based on the SARS-CoV-2 spike glycoprotein[26]. Along with the structural proteins, utilizing the non-structural and accessory proteins for the vaccine development can aid in better development of an efficacious vaccine for long term by neutralizing the mutation rate of this RNA virus. In this study, we explored the whole proteome of SARS-CoV-2 to scrutinize the highly conserved antigenic epitopes for the construction of a multi-epitope vaccine candidate that can effectively elicit both humoral and cellular mediated immune response against COVID-19. The constructed vaccine product has high population coverage and along with adaptive immunity, can as well lead to initiation of innate immune response further enhancing the generation of memory immunity.

With the continuous transmission of the virus across borders and increasing health burden on the global scale, SARS-CoV-2 demands an urgent immunization therapy. As depicted by Shan Lu, the Community Acquired Coronavirus Infection (CACI) caused by this virus can shatter the socio-economic condition worldwide and the development of vaccine against the SARS-CoV-2 should be encouraged to manage the present situation as well as it can serve as a prototype for other coronaviruses[30, 31]. Thereby, our analysis provides a platform for the development of a protective vaccine candidate that can be tested in-vitro and in-vivo and may lead to faster development of an efficient vaccine against the SARS-CoV-2 infection.

## MATERIAL AND METHOD

### 1. Retrieval of SARS-CoV-2 proteome

The complete proteome of latest reported novel Wuhan strain of SARS Coronavirus (SARS-CoV-2) was downloaded from the Nucleotide database available at National Center for Biotechnology Information (NCBI). The database was also thoroughly searched for the genomes of other available human infecting strains of Coronavirus till date and was downloaded for further analysis.

### 2. Antigenicity prediction in the Coronavirus proteome

For the antigenic analysis of the proteome of the COVID-19 strain, VaxiJen v2.0 server available at http://www.ddg-pharmfac.net/vaxijen/VaxiJen/VaxiJen.html [32]. For antigenicity prediction of the Proteins of SARS-CoV-2 with higher accuracy, Virus model available at the VaxiJen server with a threshold of 0.4 was utilized. Proteins with a VaxiJen score ≥ 0.4 were taken for epitope prediction analysis.

### 3. Selection of MHC I and MHC II alleles

MHC class I and II alleles were selected on the basis of their occurrence worldwide. We focused specially on the countries which are severely affected by the deadly SARS-CoV-2 strain. For MHC class I, following 14 HLA alleles HLA-A01:01, HLA-A02:01, HLA-A02:03, HLA-A02:06, HLA-A03:01, HLA-A11:01, HLA-A24:02, HLA-A31:01, HLA-A33:01,HLA-B07:02, HLA-B35:01, HLA-B51:01, HLA-B54:01, HLA-B58:01 were used that cover 89.84 % of the total world population as predicted by The Allele frequency Net Database [33]. For MHC II, DRB1_0101, DRB1_0301, DRB1_0401, DRB1_0405, DRB1_0701, DRB1_0802, DRB1_0901, DRB1_1101, DRB1_1201, DRB1_1302, DRB1_1501, HLA-DPA10103-DPB10101, HLA-DPA10201-DPB10201, HLA-DPA10301-DPB10301, HLA-DQA10101-DQB10201, HLA-DQA10301-DQB10301, HLA-DQA10401-DQB10401, HLA-DQA10501-DQB10501, HLA-DQA10102-DQB10202, HLA-DQA10402-DQB10402 covering 99.71 % world population were used for the epitope screening.

### 4. Helper T cell epitope prediction

Various online servers were explored for the accurate prediction of Helper T Lymphocytes (HTL) epitopes. Firstly, the antigenic proteins were analyzed by using NetMHCIIPan 3.2 server (http://www.cbs.dtu.dk/services/NetMHCIIpan/) [34]. The server provides the epitope predictions for three human MHC class II isotypes that includes HLA-DR, HLA-DP and HLA-DQ. We explored NetMHCIIPan server for HTL epitope predictions for the selected 20 MHC class II alleles. The length for the HTL epitopes was kept 15 mer. The server predicts the binding affinity of the epitopes with the respective HLA allele and provides an IC_50_ (in nanoMolar)and %Rank for each MHC class II allele – epitope pair which can be utilized for initial screening of the potential HTL epitopes. The IC_50_ and %Rank were then utilized to screen the probable strong HTL binders from the weak binders and non-binders.

The predicted strong binders for each HLA allele were further analyzed by MHCII binding prediction tool available at IEDB server (http://tools.immuneepitope.org/mhcii/) [35] and only the epitopes with rank ≤1 were taken for further analysis. Thereafter, the selected epitopes were checked for their antigenicity by using VaxiJen server, allergenicity by AlgPred (https://webs.iiitd.edu.in/raghava/algpred/submission.html) [36] and toxicity by ToxinPred (https://webs.iiitd.edu.in/raghava/toxinpred/motif_scan.php) [37]. Only the epitopes with VaxiJen score ≥ 0.4 and predicted to be non-toxic and non-allergens, were taken for further screening. The best HLA-non-allergenic epitope pair with the highest VaxiJen score was taken for constructing the multi-epitope vaccine.

### 5. Cytotoxic T cell epitope prediction

For Cytotoxic T cell (CTL) epitope prediction, NetMHCPan 4.0 [38]based on artificial neural network (ANN) was utilized for the 14 selected HLA class I molecules. The predicted strong binders were then checked for their antigenicity by using VaxiJen server. Further, immunogenicity of the epitopes was checked using class I Immunogenicity tool available at IEDB server (http://tools.iedb.org/immunogenicity/) [39] and for the allergenicity, the AllerTop v. 2.0 was used (https://www.ddg-pharmfac.net/AllerTOP/method.html) [40]. Thereafter, the non-allergenic strong binders with a positive immunogenicity score were checked for their toxic nature by utilizing the ToxinPred server (https://webs.iiitd.edu.in/raghava/toxinpred/motif_scan.php).

### 6. Epitope Conservation analysis

The presence of selected best epitopes in all the reported human infecting SARS-coronavirus strain were checked by using Epitope Conservancy Analysis tool of IEDB (http://tools.iedb.org/conservancy/) [41]. Only the epitopes having 100% conservation were used for multi-epitope vaccine construction.

### 7. Molecular interaction of the HLA-epitope pair

The MHC class I ad class II molecules were downloaded from RCSB PDB database (https://www.rcsb.org/). Those that were not available were in PDB database were retrieved from pHLA database (https://www.phla3d.com.br/) [42]. The structures of the SARS-CoV-2 proteins were constructed using I-TASSER (https://zhanglab.ccmb.med.umich.edu/I-TASSER/) [43]and HTL and CTL epitopes were mapped and their structures were retrieved using PyMol tool. For the molecular interaction analysis of the predicted best HLA-epitope pairs for both MHC class I and class II alleles, ClusPro protein-protein docking tool (https://cluspro.org/login.php) [44] was utilized.

### 8. B-Cell epitope prediction

ABCpred tool based on artificial neural network was explored for B-cell epitope prediction (https://webs.iiitd.edu.in/raghava/abcpred/index.html) [45]. For higher accuracy, the threshold for the prediction was kept 0.90. The predicted epitopes were checked for their antigenicity using VaxiJen server, allergenicity by using AlgPred and toxicity by ToxinPred server.

### 9. Designing of Vaccine construct

For constructing a multi-epitope vaccine construct, the selected best HTL, CTL and B-cell epitopes were joined by using GPGPG, AAG and KK linkers respectively. For the better immunogenic response, four adjuvants namely, β-defensin, universal memory T cell helper peptide (TpD), PADRE sequence and a M cell ligand were added by using EAAAK linker into the vaccine construct.

### 10. Antigenicity and allergenicity analysis of the vaccine construct

The antigenicity of the vaccine construct, VaxiJen server was exploited. For the allergenicity analysis of the vaccine construct, three different tools namely AlgPred, AllerTop and AllergenFP v.1.0 (http://ddg-pharmfac.net/AllergenFP/index.html) [46]were utilized.

### 11. Population coverage of the vaccine construct

To check the population coverage of the vaccine construct, Population coverage tool available at IEDB (http://tools.iedb.org/population/) [47] server was utilized. The HLA class I and class II alleles in the final construct were entered in the tool and the population coverage of the alleles were calculated for the top 26 countries that are severely affected by the SARS-COV-2 virus. The analysis was performed for HLA I and HLA II separately as well as in combination.

### 12. Physiochemical property analysis of the multi-epitope vaccine construct

Expasy’s ProtParam (https://web.expasy.org/protparam/) [48] was explored for the physiochemical properties evaluation of the vaccine construct. ProtParam evaluates the peptide sequence and provides the molecular weight, theoretical pI, extinction coefficient, estimated half-life, instability index, and grand average of hydropathicity (GRAVY). Solubility of the construct was calculated using SolPro tool available at SCRATCH protein predictor server (http://scratch.proteomics.ics.uci.edu/) [49].

### 13. Vaccine construct structure prediction and validation

Secondary structure of the multi-epitope vaccine construct was predicted using SOPMA(https://npsa-prabi.ibcp.fr/cgibin/npsa_automat.pl?page=/NPSA/npsa_sopma.html) [50] and PSI-PRED (http://bioinf.cs.ucl.ac.uk/psipred/) server [51]. For the tertiary structure prediction of the vaccine construct Robetta server (http://robetta.bakerlab.org/) based on ab initio and homology modelling was utilized [52]. The predicted structure was refined by using 3D Refine (http://sysbio.rnet.missouri.edu/3Drefine/) [53]and further by GalaxyRefine (http://galaxy.seoklab.org/cgi-bin/submit.cgi?type=REFINE) [54]. The structures were evaluated by constructing Ramachandran plot using RAMPAGE (http://mordred.bioc.cam.ac.uk/~rapper/rampage.php) and the quality was assessed using ERRAT server (https://servicesn.mbi.ucla.edu/ERRAT/).

### 14. Standard molecular dynamics of the vaccine construct

The refined modelled structure of the multi-epitope vaccine construct was further evaluated for its stability in the real environment by simulating it in a water sphere using NAMD-standard molecular dynamics tool (https://www.ks.uiuc.edu/Research/namd/) by using parallel processors. The required structure files (.psf) were generated by psfgen using Visual Molecular Dynamics (VMD) tool v.1.9.3 by utlizing CHARMM force fields for proteins. Initially, a 10,000 steps energy minimization was performed followed by subsequent heating the system from 0 K to 310 K. Thereafter, a 20 ns standard molecular dynamics was performed and trajectory DCD file generated was used to evaluate RMSD. The change in kinetic, potential and total energy was evaluated for these 20 ns simulation using VMD.

### 15. Interaction analysis of Vaccine construct with immune system molecules and there MD analysis

To check the interaction of the multi-epitope vaccine construct with two immunoreceptors, TLR-3 and TLR-8, ClusPro docking server was used and the resultant best complexes were then simulated for 20 ns in a water sphere using NAMD.

### 16. Codon optimization and *In silico* cloning

For the expression and isolation of the constructed multi-epitope vaccine in *Escherichia coli* K12 satrain, the construct was first converted to cDNA using Reverse translate tool available at Expasy server. The resultant DNA was further optimized for enhanced protein expression by using JCAT server [55]. Finally, the cDNA construct was inserted into the pET28a (+) vector using *HindIII* and *BamHI* restriction sites.

## RESULTS

### 1. Retrieval of structural and non-structural proteins of SARS-CoV-2 and their antigenicity analysis

The complete proteome of the Wuhan seafood market pneumonia virus isolate Wuhan-Hu-1 (nucleotide accession number - NC_045512.2) was retrieved from the NCBI database. SARS-CoV-2 proteome consists of a polyprotein ORF1ab and several structural proteins. ORF1ab encodes for various non-structural proteins including Host translation inhibitor nsp1, RNA dependent RNA polymerase (RnRp), Helicase, Guanine-N7 methyltransferase etc. that plays a critical role in the virus multiplication and survival inside the host. Structural proteins include Spike glycoprotein, Envelope protein, membrane protein and nucleocapsid protein. The Fasta sequences of all these proteins were retrieved from NCBI and used for epitope screening (Table 1). All the sequences were analyzed for their antigenic nature by using VaxiJen server which is based upon auto cross covariance (ACC) transformation. VaxiJen classifies the proteins into antigens and non-antigens solely based on the physiochemical properties and is sequence alignment independent. The ACC score threshold for the virus model is kept 0.4 that increases the prediction accuracy. All the SARS-CoV-2 proteins except NSP16 (2’-O-ribose methyltransferase) resulted in the ACC score higher than 0.4 depicting the antigenic nature of the viral proteome (Table 1). All the antigenic proteins were further screened for the presence of HTL, CTL and B-cell epitopes by using various tools and databases (Figure 1).

**Table 1.** List of SARS-CoV proteins used for antigenicity prediction using VaxiJen server. The antigenic proteins with Antigenicity Score ≥0.4 were taken for epitope screening.

| NCBI accession number | SARS-CoV-2 proteins |  | Antigenicity score | Antigenicity prediction |
| --- | --- | --- | --- | --- |
| QHD43415_1 | ORF1ab | Host translation inhibitor (nsp1) | 0.4064 | Probable ANTIGEN |
| QHD43415_2 |  | Non-structural protein 2 (nsp2) | 0.4034 | Probable ANTIGEN |
| QHD43415_3 |  | Papain-like proteinase | 0.5142 | Probable ANTIGEN |
| QHD43415_4 |  | Non-structural protein 4 (nsp4) | 0.4575 | Probable ANTIGEN |
| QHD43415_5 |  | Proteinase | 0.4159 | Probable ANTIGEN |
| QHD43415_6 |  | Non-structural protein 6 (nsp6) | 0.5813 | Probable ANTIGEN |
| QHD43415_7 |  | Non-structural protein 7 (nsp7) | 0.4167 | Probable ANTIGEN |
| QHD43415_8 |  | Non-structural protein 8 (nsp8) | 0.4008 | Probable ANTIGEN |
| QHD43415_9 |  | Non-structural protein 9 (nsp9) | 0.6476 | Probable ANTIGEN |
| QHD43415_1 |  | Non-structural protein 10 (nsp10) | 0.4039 | Probable ANTIGEN |
| QHD43415_1 |  | RNA-directed RNA polymerase (RdRp) | 0.4064 | Probable ANTIGEN |
| QHD43415_1 |  | Helicase (Hel) | 0.448 | Probable ANTIGEN |
| QHD43415_1 |  | Guanine-N7 methyltransferase (ExoN) | 0.4138 | Probable ANTIGEN |
| QHD43415_1 |  | Uridylate-specific endoribonuclease (NendoU) | 0.5554 | Probable ANTIGEN |
| QHD43415_1 |  | 2'-O-methyltransferase (2'-O-MT) | 0.38 | Probable NON-ANTIGEN |
| QHD43416 |  | Surface glycoprotein | 0.4646 | Probable ANTIGEN |
| QHD43417 |  | ORF3a | 0.4945 | Probable ANTIGEN |
| QHD43418 |  | ORF4 (E Protein) | 0.6025 | Probable ANTIGEN |
| QHD43419 |  | Membrane Protein | 0.5102 | Probable ANTIGEN |
| QHD43420 |  | ORF6 | 0.6131 | Probable ANTIGEN |
| QHD43421 |  | ORF7a | 0.6441 | Probable ANTIGEN |
| QHD43422 |  | ORF8 | 0.6502 | Probable ANTIGEN |
| QHD43423 |  | Nucleocapsid Protein | 0.5059 | Probable ANTIGEN |
| QHI42199 |  | ORF10 | 0.7185 | Probable ANTIGEN |

**Figure 1.**
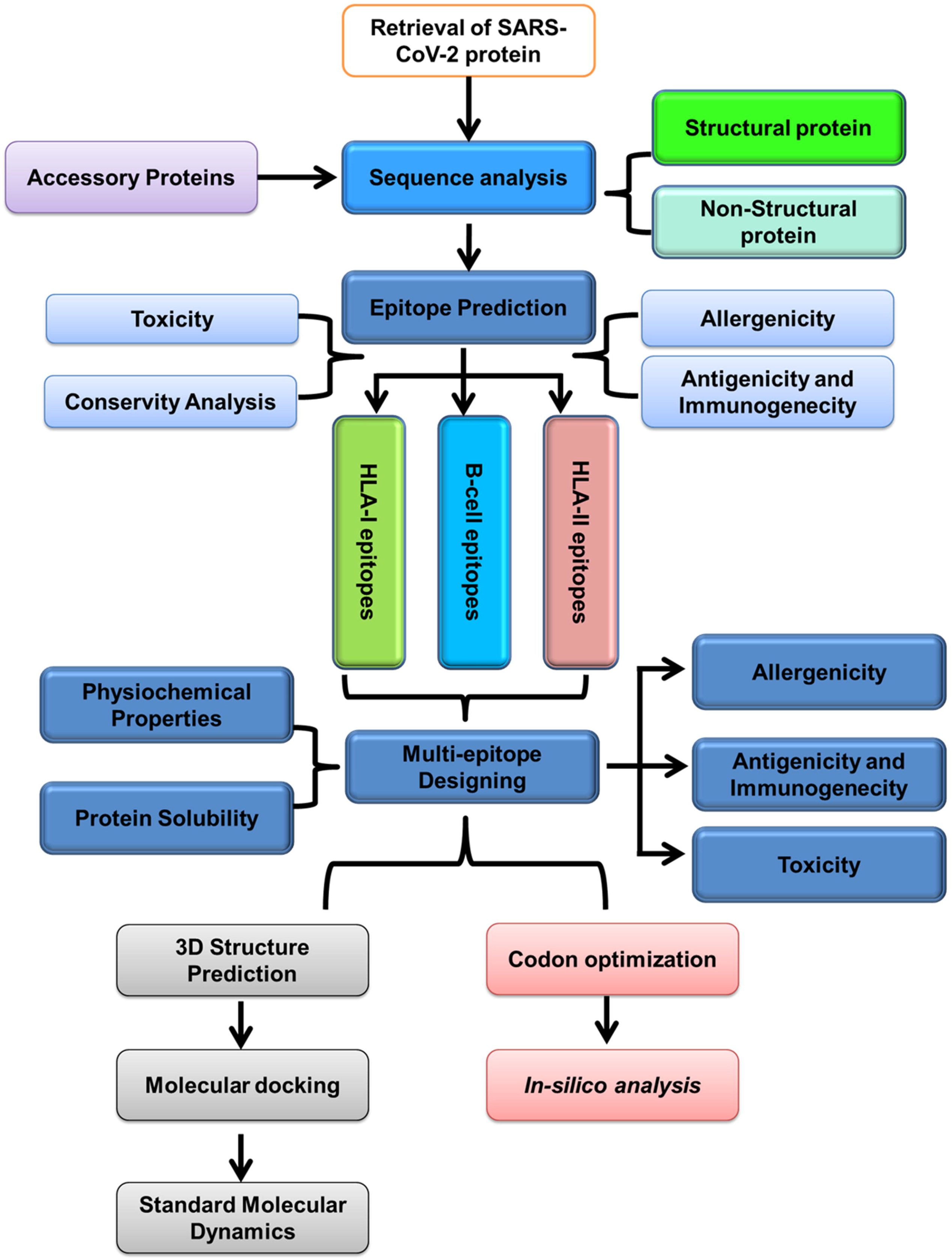
Schematic representation of next generation vaccinology approach used for the prediction of Multi-epitope vaccine construct for SARS-CoV-2.

### 2. Epitope screening revealed the presence of high affinity HTL, CTL and B-cell epitopes in the antigenic proteome of SARS-CoV-2

T-Lymphocytes play a central role in activating the cell mediated innate and adaptive immune response against the foreign particles. As well they are the sole players in generating immunological memory that provides long lasting immune response. Thus, the vaccinology approach revolves around the screening of proteome for the high affinity Helper T Lymphocytes (HTL) and Cytotoxic T Lymphocytes (CTLs) epitopes that can activate T_H_ and T_C_ cells. The epitopes for the activation of HTLs and CTLs are processed and presented by various human leukocyte antigen (HLA) class I and class II molecules. The HLA-epitope binding steps are crucial and have high specificity and are thus considered as one of the main focus for vaccine designing. HLA shares large diversity due to the presence of several alleles in the human genome but their occurrence is not uniform world-wide. Few of the alleles are present throughout the world while some are restricted to only specific regions. Recent reports and literature survey lead us to identify the massively affected regions by COVID-19, namely China, Italy, Spain, Germany, USA, Iran, France, South Korea, Switzerland, UK, Netherlands,Austria,Belgium,Norway,Sweden,Denmark,Canada,Malaysia,Australia, Portugal, and Japan. The presence of various epitopes with different HLA-binding specificities leads to greater population coverage as the frequency of HLA allele’s expression varies with the varying ethnicities worldwide. Henceforth, fourteen HLA class I and Twenty class II alleles were selected on the basis of their occurrence in these countries. Individually, HLA class I covers 89.84 % world population, while HLA class II covers 99.71 %.

HTLs act as the central mediators of immune response and coordinates with B lymphocytes, T_C_ cells and macrophages via signaling through interleukins and cytokines synthesis. HTLs recognize the epitopes presented on MHCII biding cleft. Herein, NetMHCPanII server was utilized for predicting 15 mer HTL epitopes having high binding affinity to the selected 20 MHC II alleles. Only the strong binders for the respective HLA alleles were selected with a percentile rank of ≤1. The selected epitopes were further analyzed using another MHC II prediction tool available at IEDB server and filtered on the basis of percentile rank. The resultant epitopes were checked for their antigenicity, allergenicity and toxicity. Finally, the best HLA-epitope pairs with the highest antigenicity and were non-allergen and non-toxic were used for multi-epitope vaccine construct. On analysis, we received, 2 best HLA-epitope pairs each for Helicase and RdRp. Three best HLA-epitope pairs were obtained for non-structural protein 2 (NSP2) and Spike glycoprotein (S) and one pair each for membrane glycoprotein (M), ORF3a and ORF8 proteins. In summary, non-toxic, non-allergenic epitopes with high binding affinity and antigenicity were obtained for 15 MHC class II alleles and their respective epitopes were used for multi-epitope vaccine construct. Best binding epitopes for the remaining 5 MHC II alleles were either non-antigenic, or were allergens and thus were excluded (Supplementary Table S1).

CTLs eliminate foreign particles thereby helping in pathogen clearance and are required for maintaining the cellular integrity. MHC class I molecules represent the epitopes to the CTLs which after activation perform cytotoxic activities. Here, we used NetMHCpan v.4 for the mining of CTL epitopes that have high binding affinity with the 14 class I molecules. The strong binders obtained from NetMHCpan server were then analyzed for their immunogenicity by using class I Immunogenicity tool. Depending upon the amino acid composition, properties and their position in the epitope, class I immunogenicity tool predicts the immunogenicity of a MHC class I – epitope complex. Higher the score depicts higher immunogenicity and vice-versa. Here, all the epitopes with positive score were taken and analyzed for antigenicity, allergenicity and toxicity. Finally, the epitopes with highest antigenicity,non-allergen and non-toxic wereselectedfor the vaccine construction.Upon analysis, 3 HLA-epitope pairs were obtained for ORF3a protein, 2 pairs for membrane protein, nucleo-capsid protein, and spike glycoprotein each while one HLA-epitope pair each for NSP2, NSP4, Envelope protein, and ORF8 (Supplementary Table S2). Overlapping sequences were merged into one or both MHC class I and class II binding epitopes

Apart from cellular mediated immunity, the humoral immune response mediates pathogen clearance in the antibody dependent manner. Hence the proteome of SARC-CoV-2 was further scanned for linear B-cell epitopes by using ABCPred server. For higher selectivity and sensitivity, the threshold of ABCPred was kept 0.9. The predicted epitopes were checked for their antigenicity, allergenicity and toxicity and the best epitopes were taken for further consideration. On the basis of above criteria, we received one B-cell epitope each for Nucleocapsid, Guanine-N7 methyltransferase (ExoN), ORF3a, ORF7a, and Surface glycoprotein (Supplementary Table S3). The locations of the selected epitopes in the respective protein structures are represented by Figure 2. Overall, the antigenic 13 HTL and 12 CTL epitopes having highest affinity for the respective HLA alleles and 5 B-cell epitopes that are non-allergenic, non-toxic and can generate a potential immune response were selected for incorporation into the multi-epitope vaccine construct.

**Figure 2.**
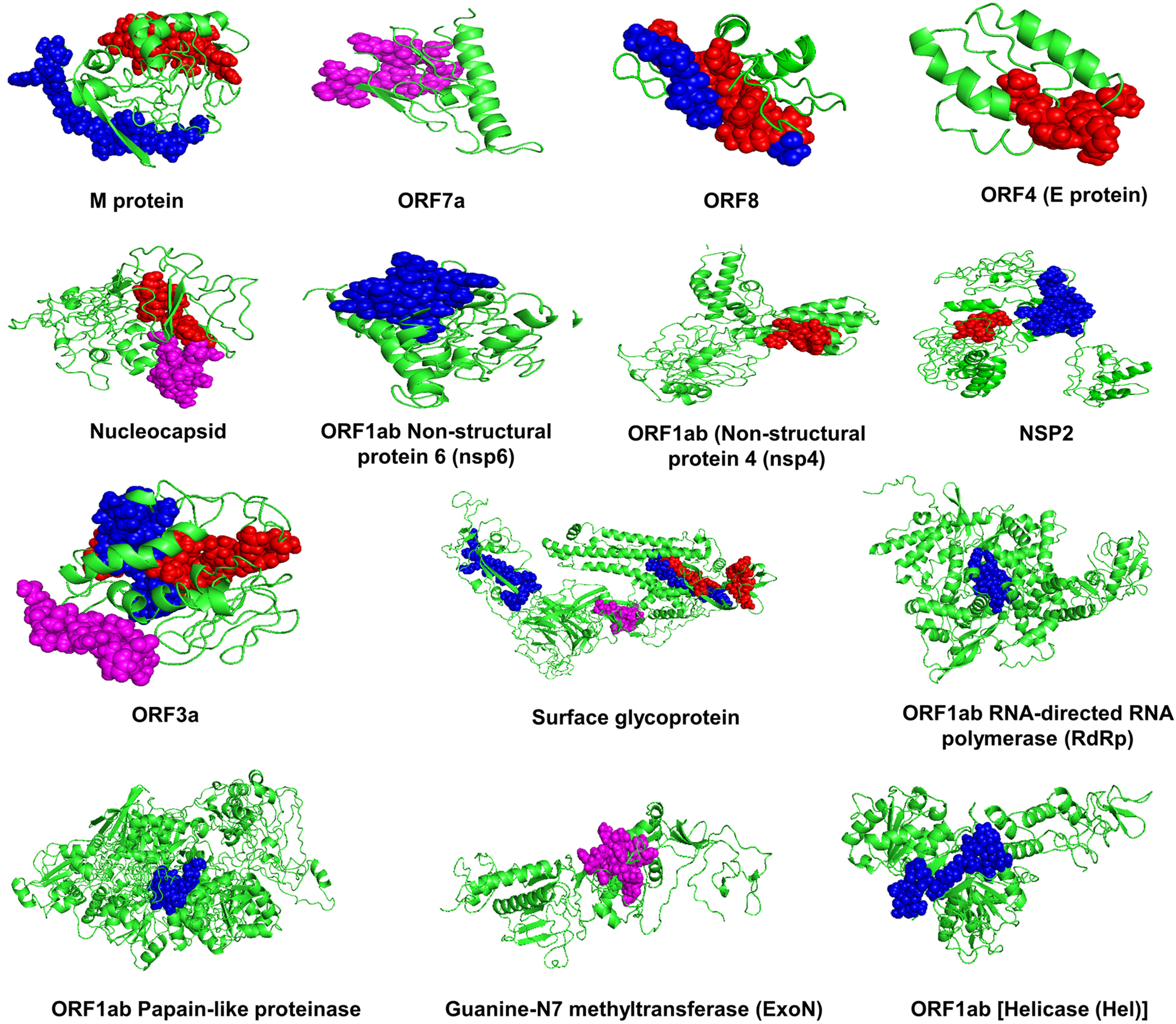
HTL, CTL and BCL epitopes in SARS-CoV-2 proteome. Location of selected Helper T-Lymohocytes epitopes (Blue), Cytotoxic T-Lymphcytes epitopes (Red) and B Cell Lymphocytes epitopes in the three dimensional structures of the antigenic proteins of SARS-CoV-2.

### 3. The screened epitopes shared high conservancy within the Coronovirus family

The presence of conserved epitopes in a vaccine can lead to an effective immunization against all the strains of the pathogen. Thus, the selected HTL, CTL and BCL epitopes were analyzed for their conservancy among the various human infecting strains of coronavirus. Interestingly, all the selected epitopes were 100 % conserved throughout the coronavirus family and this can lead to a robust vaccine development.

### 4. Molecular interaction analysis depicts the strong interaction of T-cell epitopes and HLA molecules

MHC class I and class II alleles have a specific binding groove where the peptide binds and are presented on the cell surface for immune system activation. The protein structures of SAR-CoV-2 were modelled using I-Tasser (Supplementary Figure S1). To check the molecular interaction of our selected CTL and HTL epitopes with their respective HLA alleles, we performed protein-protein docking using ClusPro. HLA alleles were taken as receptors and epitopes as ligands. ClusPro performs the interaction analysis by first generating billions of conformers eventually leading to rigid target based molecular docking. The resultant docked complexes are then clustered based on RMSD and lowest energy finally leading to energy minimization. The cartoon representation of molecular interaction between HLA alleles and epitopes are shown in Figure xx. Interaction analysis revealed favorable interaction of both CTL and HTL epitopes with the HLA allele’s epitope binding grooves. These epitopes will be presented to the T_C_ and T_H_ cells and would lead to immune response against the vaccine. (Supplementary Figure S2)

### 5. Population coverage analysis shows the broad range spectrum of the constructed vaccine

Population coverage of the vaccines depends upon the interaction of the epitopes with the number of HLA alleles that are then represented to T_C_ and T_H_ cells that elicit the immune response. Depending upon the ethnicity, the expression of HLA alleles varies throughout the world. Thus to check the population coverage of the vaccine construct, presence of selected HLA alleles were analyzed in the 26 severely affected countries by COVID-19 around the world. Individually, MHC class I alleles covered on an average 91.22 % of world population while MHC class II alleles covered 99.70 % (Figure 3A & B). As our vaccine construct harbors HTL epitopes for 15 MHC class II and CTL epitopes for 14 MHC class I alleles, the occurrence of these HLA alleles depicted the population coverage of >99% throughout the world. Five countries including United States and France showed 100% population coverage while ≥95% of population was covered in rest countries except United Kingdom depicting the broad range spectrum of the constructed vaccine (Figure3C).

**Figure 3.**
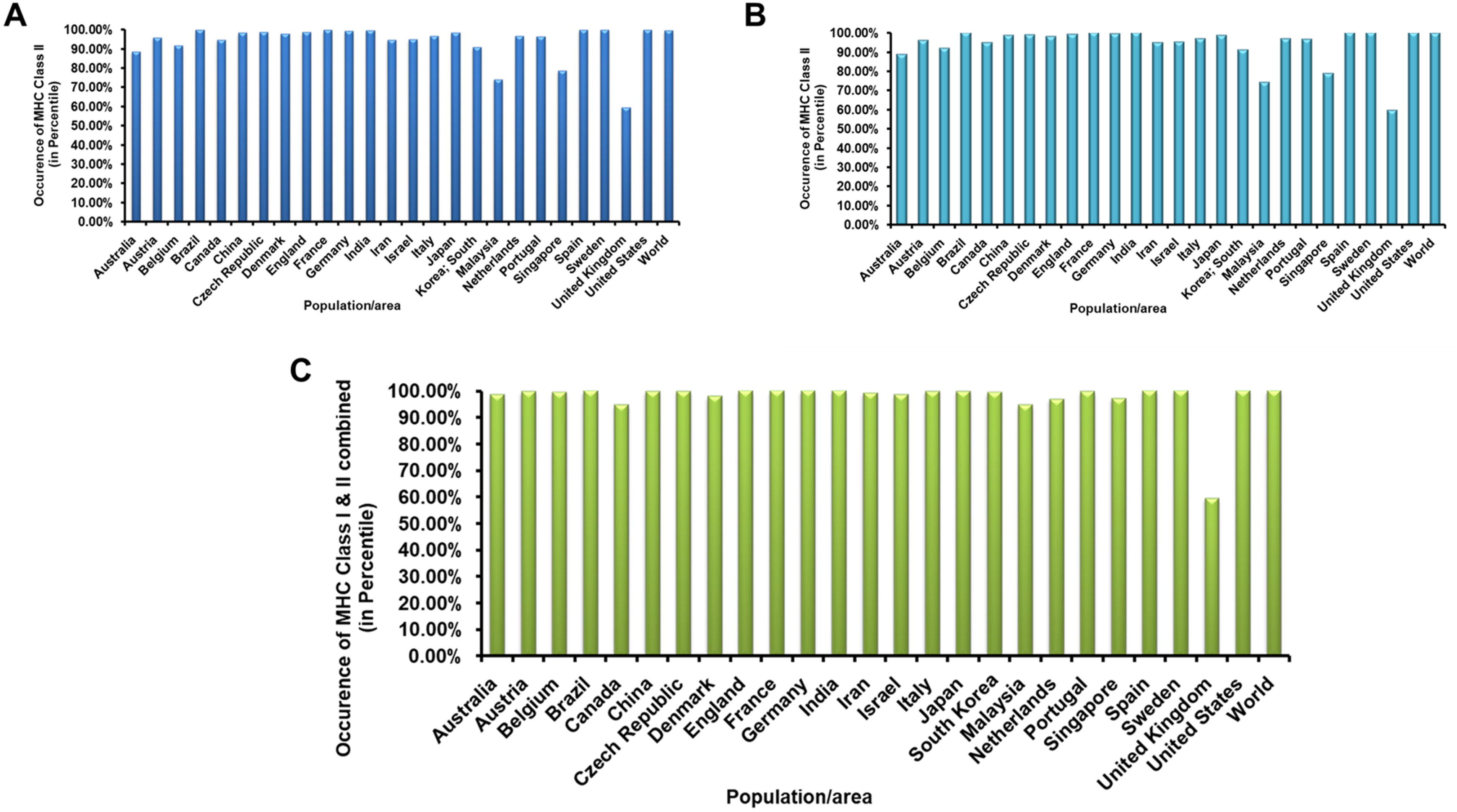
Population Coverage of the SARS-CoV-2 multi-epitope vaccine construct. (A & B) The population coverage of HLA class I and class II alleles for whom the best antigenic epitopes were screened in various countries severely affected by COVID-19 in the world. (C) Population coverages of the multi-epitope vaccine construct based on the coverages of HLA class I and class II alleles in combination in various countries.

### 6. Designing of SARS-CoV-2 multi-epitope vaccine construct

An ideal vaccine should harbor conserved epitopes, have multi-valency and can elicit both cellular and humoral mediated immune response in the host. A subunit vaccine contains minimal elements that are antigenic and required for the stimulation of prolonged protective or therapeutic immune response. In the recent times, various reports have shown the construction of multi subunit vaccine by utilizing highly antigenic HTL, CTL and BCL epitopes. Herein, we constructed a multi-epitope vaccine candidate by combining 13 HTL, 12 CTL and 4 BCL epitopes that were highly conserved, antigenic, non-toxic and non-allergens (Figure 4).

**Figure 4.**
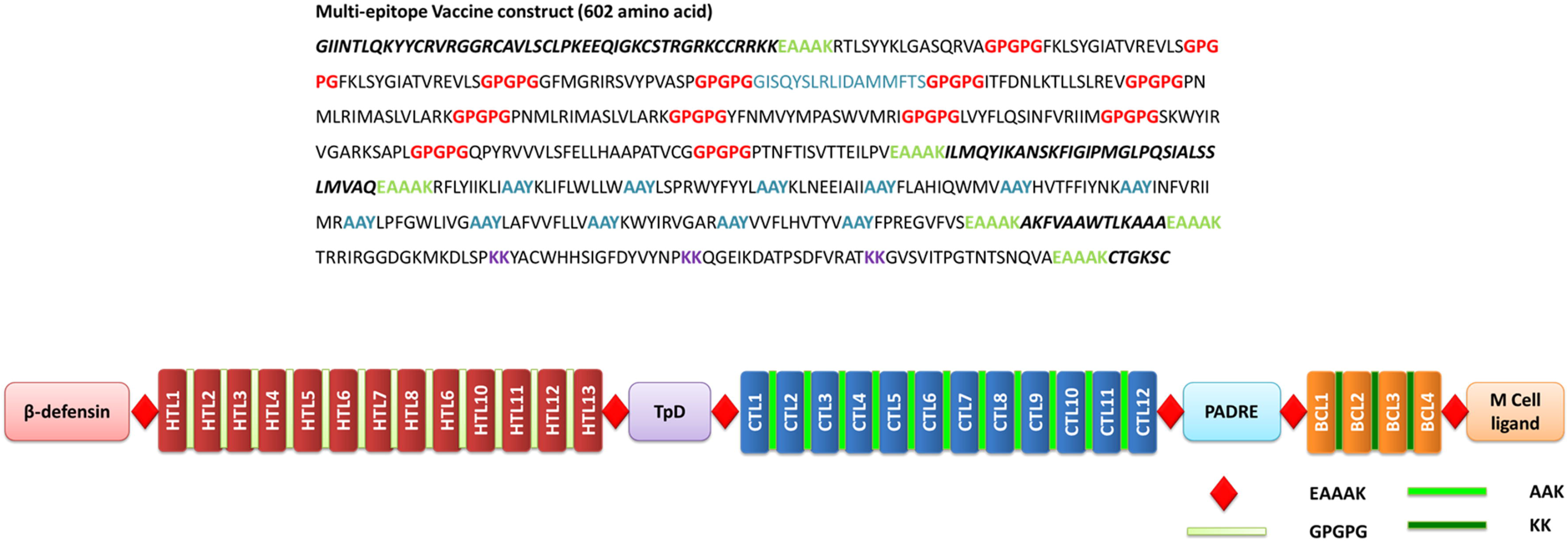
Designing SARS-CoV-2 multi-epitope vaccine construct. The Sequence of the multi-epitope vaccine construct with HTLs, CTLs and BCL epitopes along with various adjuvants and linkers. Schematic representation of the SARS-CoV-2 multi-epitope vaccine construct.

In the recent times, various small peptides are been explored that acts as an adjuvant and can potentiate the multi-epitope vaccine mediated immune response by activating humoral immunity. Taking this into consideration, apart from the conserved epitopes, four adjuvants were also added so as to boost the protective immune response against SARS-CoV-2. β-defensins are antimicrobial peptides that acts as chemoattractant for the immature dendritic cells and T-lymphocytes and induces their maturation.[56] A 45 amino acid long GIINTLQKYYCRVRGGRCAVLSCLPKEEQIGKCSTRGRKCCRRKK is well reported for its anti-microbial activity and was added at the vaccine construct. CD_4+_ T cells are the key regulators of humoral immunity as they assist B-cells in effective antibody class switching and their affinity maturation. Universal memory T helper cell peptide (TpD) is a chimeric peptide derived from tetanus and diphtheria toxoids and has a high promiscuity to interact with wide range of MHC class II alleles. Presence of a TpD in the vaccine construct leads to an effective T_h_ cell activation and enhances the immunogenicity of the vaccine. Similarly, a universal Pan DR epitope (PADRE) has been reported to enhance potency of long multi-epitope by activating CD4+ T helper cells and helps in generating high titer IgG antibodies.[57] Another small peptide CTGKSC act as an M cell ligand ad enhances the adsorption of oral vaccines from the intestinal membrane barrier.[58] Hence for enhanced immune response and transportation, these four adjuvants were added to the multi-epitope vaccine construct. The adjuvants and epitopes were arranges in the following order: β-defensins – HTLs – TpD – CTLs – PADRE – BCL – CTGKSC. To preserve independent immunogenic activities, various linkers were used for combining the adjuvants and epitopes. EAAAK was used to join the adjuvants, HTL, CTL and BCL epitopes with each other as they effectively separate various domains in multi-domain proteins. GPGPG linkers were used to join HTL epitopes, while AAY linkers were used for combining CTLs. GPGPG and AAY linkers enhances the recognition of the vaccine constructs by MHC I and MHC II machineries. Similarly, KK linkers were utilized for combining B-cell epitopes. KK linker acts as target for Cathepsin-B that separates the epitopes before MHC class II mediated antigen presentation to the Th0 cells. Overall, six EAAAK, 12 GPGPG, 11 AAY and 3 KK linkers were added to the subunit vaccine making it a 602 amino acid long vaccine construct (Figure 4).

### 7. Physiochemical properties, antigenicity, allergenicity, and toxicity analysis revealed the efficacy and safety of the predicted vaccine construct

Analysis of the physiochemical properties of a multi-subunit vaccine helps in analyzing the optimal immune response of the vaccine and its stability inside the host. ProtParam was utilized for accessing the molecular weight, pI, stability and grand average of hydropathy (GRAVY) constant of the SARS-CoV-2 multi-epitope vaccine construct. Molecular weight of the vaccine construct was observed to be 65.3 kDa while pI value was 10.06. Instability index of 31.93 (<40) depicted the stable nature of the vaccine construct while the estimated half-life in mammalian reticulocytes was observed to be >30 hours. GRAVY constant of 0.213 depicted the hydrophilicity of the vaccine construct and was predicted to be soluble with a probability of `70 % as predicted by SolPro (Table 2). Antigenicity is a preliminary requisite of a successful vaccine candidate. A vaccine construct must possess both immunogenicity and antigenicity to elicit the humoral and cell-mediated immune response. Upon analyzing the vaccine construct sequence in VaxiJen server, the constructed vaccine was found to be antigenic in nature with an overall prediction score of 0.6199. The score signifies the antigenic potential of the vaccine construct that may have the potential to evoke the immune response inside the host. On allergenicity analysis, all the three tools AlgPred, AllergenFP and AllerTop supported the non-allergenic nature of the vaccine construct while toxicity analysis revealed the non-toxic behavior of the construct (Table 2). In summary, the constructed epitope was observed to be stable, soluble, antigenic, non-allergenic and non-toxic.

**Table 2.**
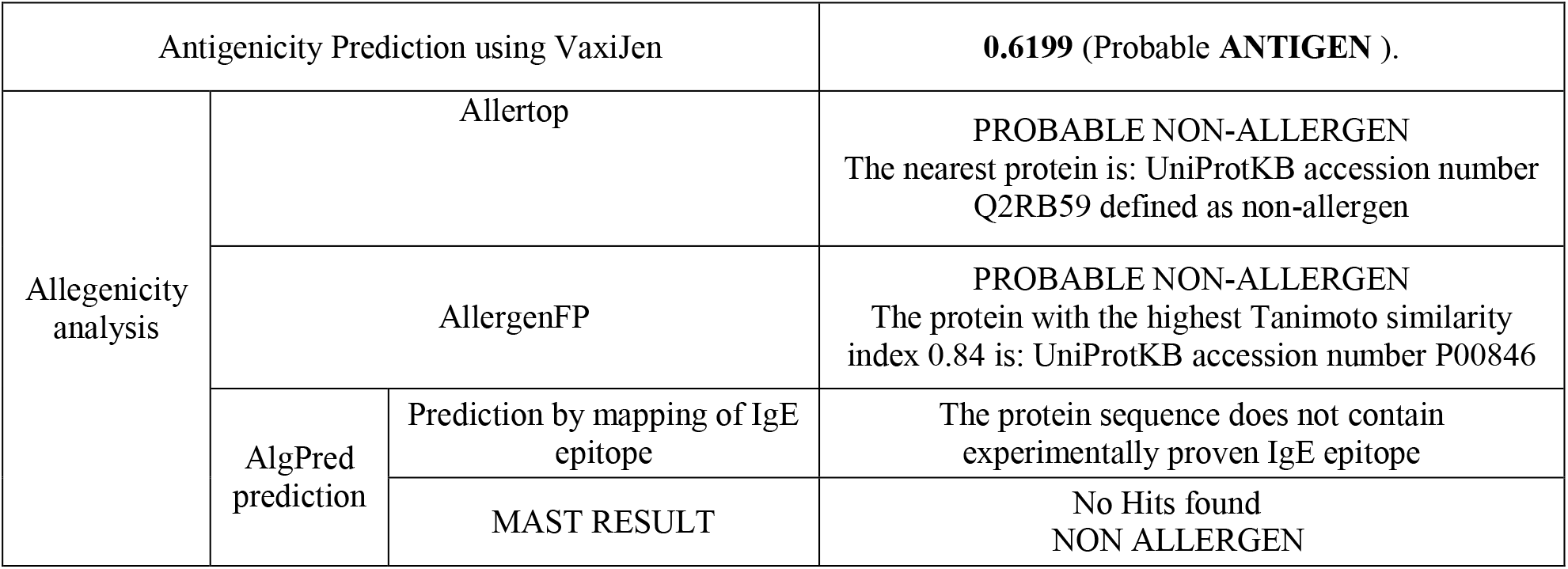
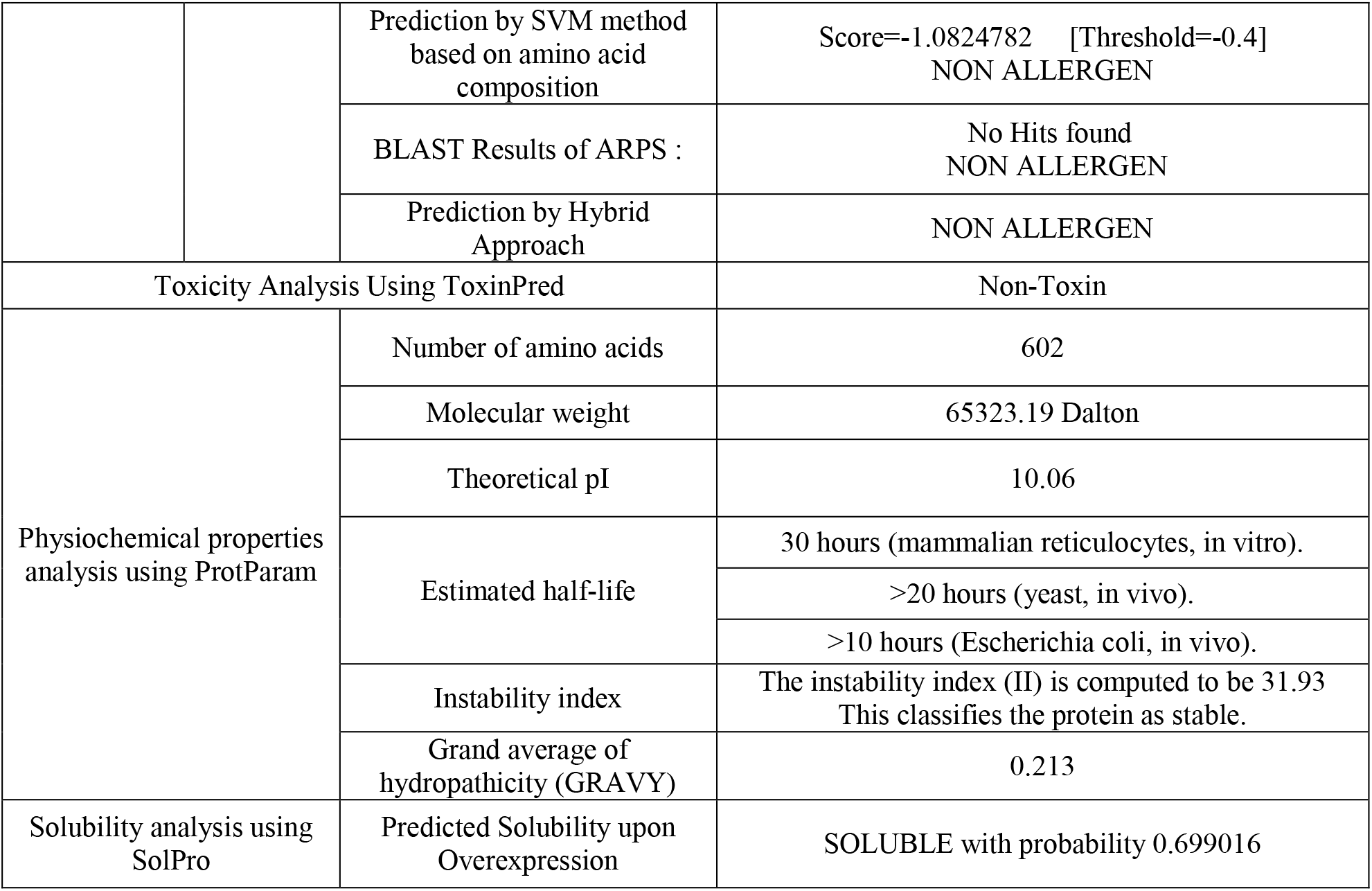
The results of physiochemical properties, antigenicity, allergenicity, and toxicity analysis of the SARS-CoV-2 multi-epitope vaccine construct.

### 8. Structure modelling and validation results in the construction of feasible 3D structure of the vaccine construct

Secondary and tertiary structure helps in the functional annotation of the multi-epitope vaccines. It also helps in analyzing the interaction of this vaccine construct with the immunological receptors like TLRs. Secondary structure of the protein was analyzed by using SOPMA and PSIPRED server that revealed the presence of ~40 % alpha helix, ~20 % β-sheet, ~30 % coils and ~6 % β turns in the vaccine construct (Figure 5 and Supplementary Figure S3). Tertiary structure of the vaccine was predicted by threading based homology modelling using Robetta server (Figure 6A). Ramachandran plot analysis of the modelled structure revealed the presence of 95.5 % residues in the most favoured regions and 4.5 % in the additionally allowed regions (Figure 6B). Quality factor was analyzed by using ERRAT2 server and was obtained to be 90.47 depicting a good modelled structure (Figure 6C). To further refine the modelled structure, 3D refine and GalaxyRefine were utilized that leads to 97.4 % residues in the most favoured region and 2.6 % residues in the additionally allowed region. The quality factor of the refined structure was observed to be 94.29% depicting the improvement of the modelled structure (Figure 6D-F).

**Figure 5.**
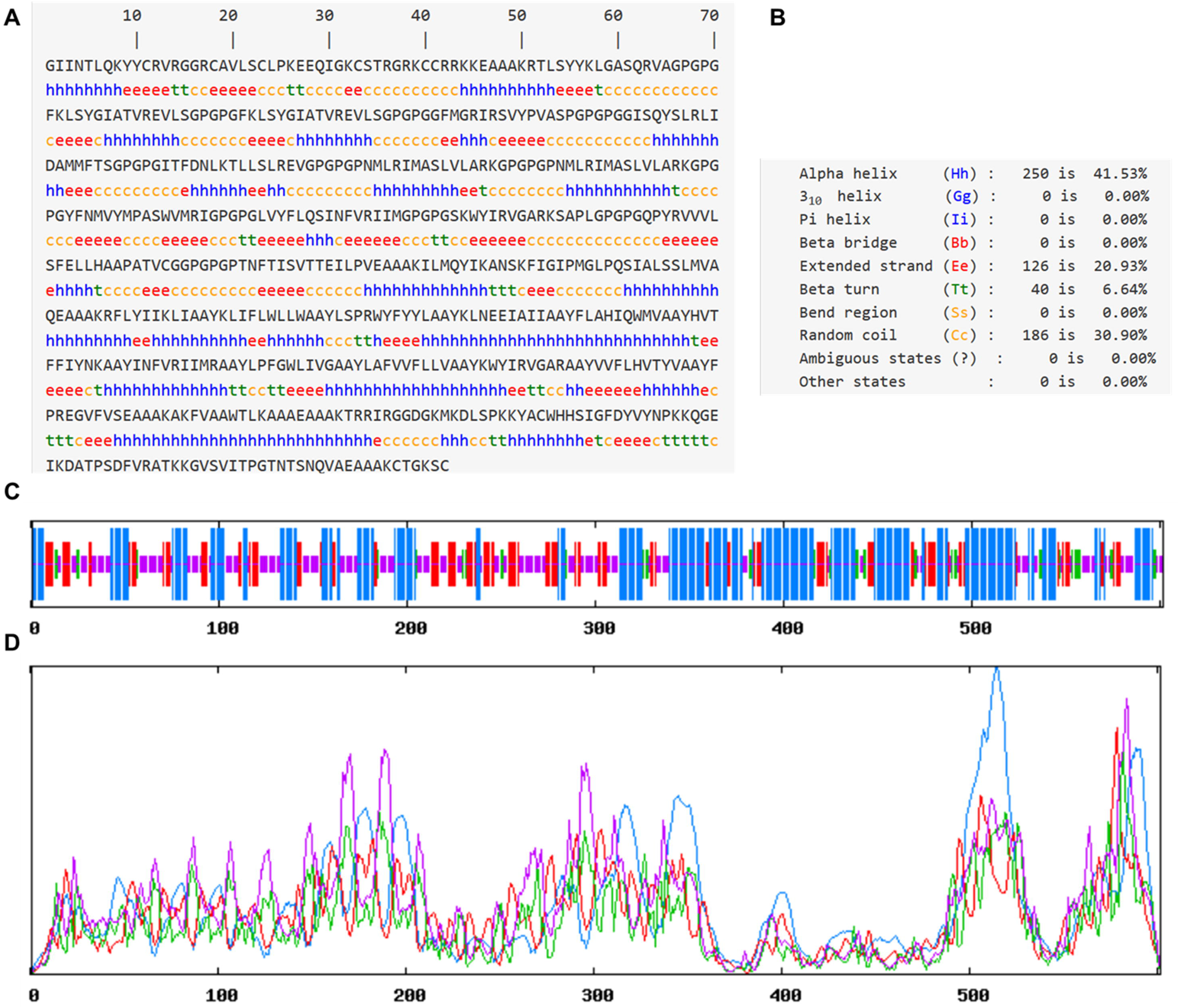
Secondary Structure prediction of SARS-CoV-2 multi-epitope vaccine construct. A. The sequence of the vaccine construct along with the predicted secondary structure. B. The overall percentage of various secondary structures in the vaccine construct as predicted by SOPMA server. C Pictorial representations of various secondary structures in the multi-epitope vaccine construct. D. Propensity of occurrence of various secondary structures according to the residues in the vaccine construct. The secondary structure at a particular residue was predicted by the height of the peak.

**Figure 6.**
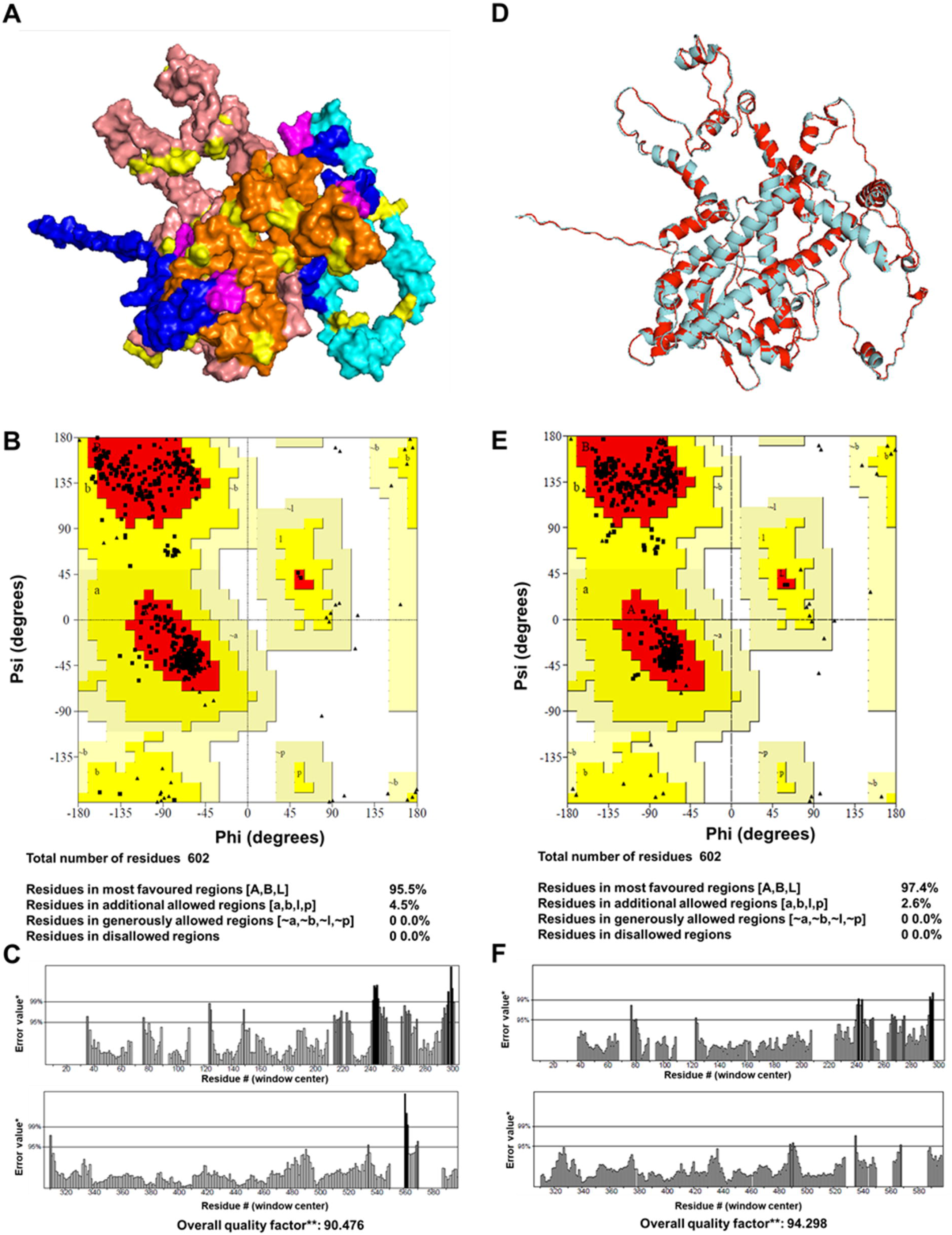
Tertiary Structure and validation of the SARS-CoV-2 multi-epitope vaccine construct. (A) Represents the modelled structure of the vaccine construct where HTL epitopes are depicted by Salmon Pink color, CTL by Orange, BCL epitopes by Cyan, and adjuvants by Blue. (B & C) Ramachandran plot and ERRAT plot generated for the modelled vaccine construct. (D) The modelled (Cyan) and refined structure (Red) of the SARS-CoV-2 multi-epitope vaccine construct depicting the changes in the structure before and after refinement. (E&F) Ramachandran plot and ERRAT plot generated for the refined structure of the vaccine construct.

### 9. Structure dynamics simulation of the vaccine construct depicts the stability of the modelled vaccine construct

Refined structure of the model construct was further check for its stability in real environment by simulating it for 20 ns in a water sphere. 10,000 steps energy minimization was performed to minimize the potential energy of the system. Unnecessary or false geometry of the protein structures are repaid by performing energy minimization resulting in a more stable stoichiometry. Before energy minimization, the potential energy was observed to be −447451.2005 Kcal/mol. After 10,000 steps the protein was minimized with a potential energy of −676040.4692 kcal/mol (Figure 7A). The system was subsequently heated from 0 K to 310 K and then 10 ns molecular dynamic simulation was performed. The temperature remained constant throughout the simulation process (Figure 7B). The DCD trajectory file was analysed for analysing the movement of the atoms during the simulation course and RMSD was calculated. Upon analysing the RMSD of SARS-CoV-2 vaccine construct, it was observed that, the system gained equilibrium at ~4 ns and then remained constant till 10 ns depicting the stability of the vaccine construct (Figure 7C). Furthermore, upon analysing the change in kinetic, potential and total energy of the system, it was observed that after a quick initial change in all the three, they remained constant throughout the simulation further strengthening the stability of the vaccine construct (Figure 7D-F). Furthermore, the bond energy, VdW energy, dihedral and improper dihedral energy analysis revealed no change throughout the 10 ns dynamics simulation. Root mean square fluctuation (RMSF) analysis revealed the rigidness of the atoms in vaccine construct with a slight mobility observed at ~ 120 and 501 residues (Supplementary Figure S4). Thus, simulation analysis revealed the stability of the vaccine construct in the real environment. The refined vaccine construct structure was therefore used for interaction analysis with immunological receptor TLR3.

**Figure 7.**
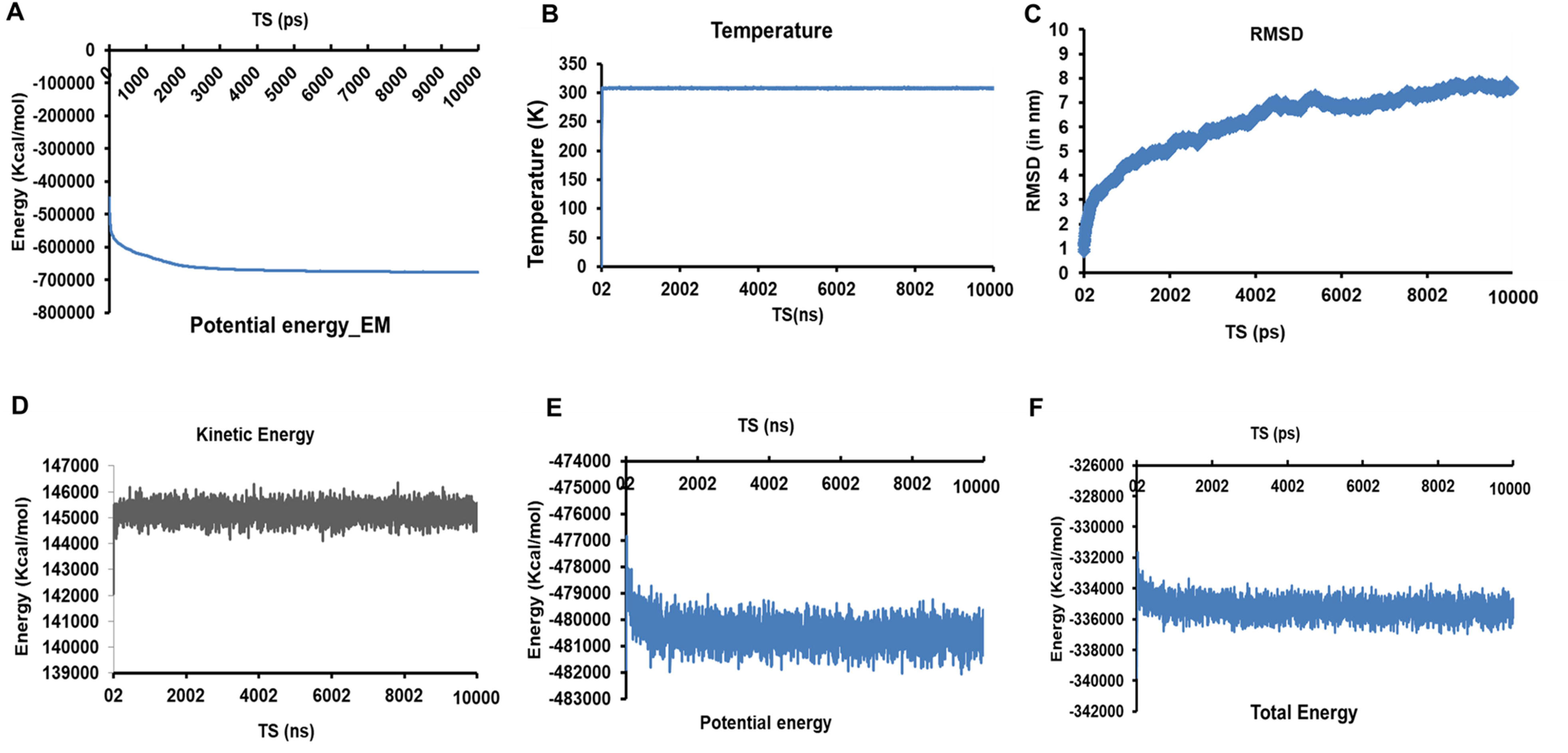
Standard molecular Dynamics analysis of SARS-CoV-2 multi-epitope vaccine construct. (A) Representation of change in potential energy vs time steps(in ns) during energy minimization of the vaccine construct. (B)Temperature Vs Time steps showing the constant temperature throughout 10 ns simulation study. (C) RMSD Vs TS depicting the root mean square deviation of the atoms of Multi-epitope vaccine construct during the 10 ns dynamic simulation. (D – F) Energy plots representing Kinetic, potential and total energy w.r.t. TS for the Vaccine construct system in a water sphere.

### 10. Interaction of vaccine construct with immunological receptor depicts the strong feasible binding

Protein – protein docking of immune receptor TLR3 and the constructed vaccine was carried out using ClusPro server. In total 30 TLR3-vaccine construct complexes were generated. The fifth model with the lowest energy weighted score of −1427.4 was considered as best TLR3-vaccine complex (Figure 8 A-B and Supplementary Table S4). The vaccine construct interacted with the ligand binding grove of TLR-3 generating a strong TLR3-vaccine construct complex. The stability of the complex was further analyzed by performing 10ns standard molecular dynamics simulation studies using NAMD suite. The 10,000 steps energy minimization of the complex lead to the generation of a minimized energy complex with energy of −756791.5601 Kcal/mol (Figure 8C). After subsequent heating the system from 0 K to 310 K, the temperature was kept constant throughout (Figure 8D). Energy plots depicted no major changes throughout the simulation (Figure 8E-G and Supplementary Figure S5). RMSF analysis of the complex showed a mobility region at ~740^th^ residue and ~1100 residue of the TLR3-vaccine complex (Figure 8H). On trajectory analysis, the RMSD plot was constructed that revealed the major deviations during the initial 1.5 ns, thereafter the system remained constant till 10ns.

**Figure 8.**
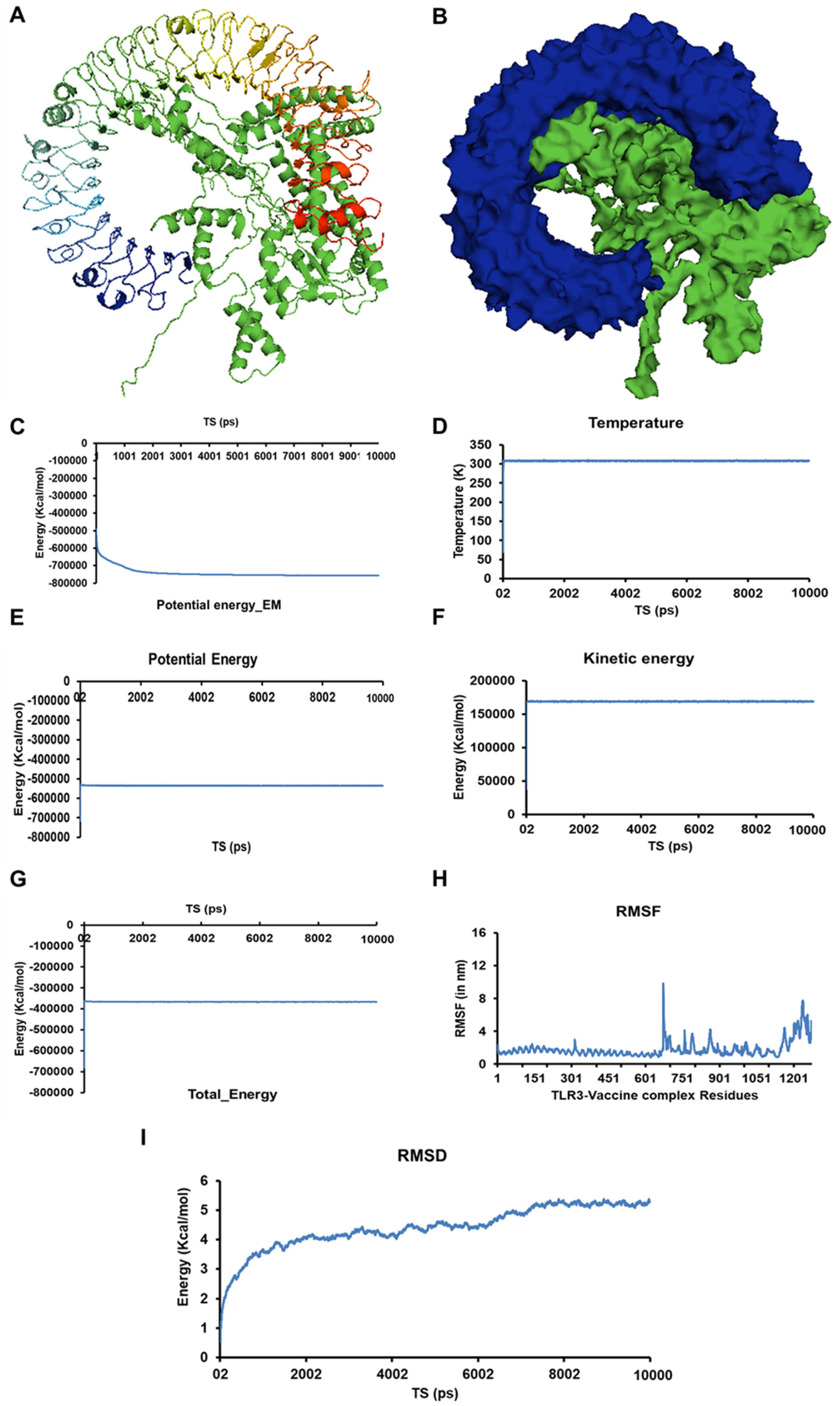
Molecular interaction analysis of SARS-CoV-2 multi-epitope vaccine construct with a immunological receptor Human TLR3. (A & B) Representation of TLR3-vaccine construct in ribbon (A) and surface form (Blue Human TLR3 ad Green-multi-epitope vaccine construct). (C) Representation of change in potential energy vs time steps (in ns) of the TLR3-vaccine construct during energy minimization. (D)Temperature Vs Time steps showing the constant temperature throughout 10 ns simulation study. (E – G) Energy plots representing Kinetic, potential and total energy w.r.t. TS for the Vaccine construct system in a water sphere. (H) RMSF Vs TS for the residues of TLR3-vaccine construct complex (I) RMSD Vs TS depicting the root mean square deviation of the atoms of TLR3-vaccine construct during the 10 ns dynamic simulation.

### 11. cDNA was optimized for the optimal expression of the vaccine product

For the insertion of the vaccine construct into a plasmid vector, the 602 amino acid long protein sequence was reverse translated to cDNA of 1806 nucleotide length. The expression system of the expression host varies with each other and the cDNA needs to be adapted as per the host codon usage. For the optimal expression of the vaccine product in *Escherichia coli K12* host, the resultant cDNA was codon optimized according using JCAT server. Also, during optimization, rho-independent transcription terminator and prokaryotic ribosomal binding sites were avoided in middle of the cDNA sequence so as to generate an optimal and complete protein expression. Further, for inserting the construct in the cloning vector, the cleavage sites of *BamHI* and *HindIII* were also avoided. The CAI value (codon adaptation index) of the cDNA before adaptation was observed to be 0.5379 and a GC content of 59.52 % (Figure 9A). After adaptation, the CAI score of the improved sequence was increased to 0.946 with 52.54 % GC content (Figure 9B and Supplementary data S1). The enhanced CAI score depicts the presence of most abundant codons in *Escherichia coli* K12. The adapted cDNA sequence was used for *In silico* cloning purpose.

**Figure 9.**
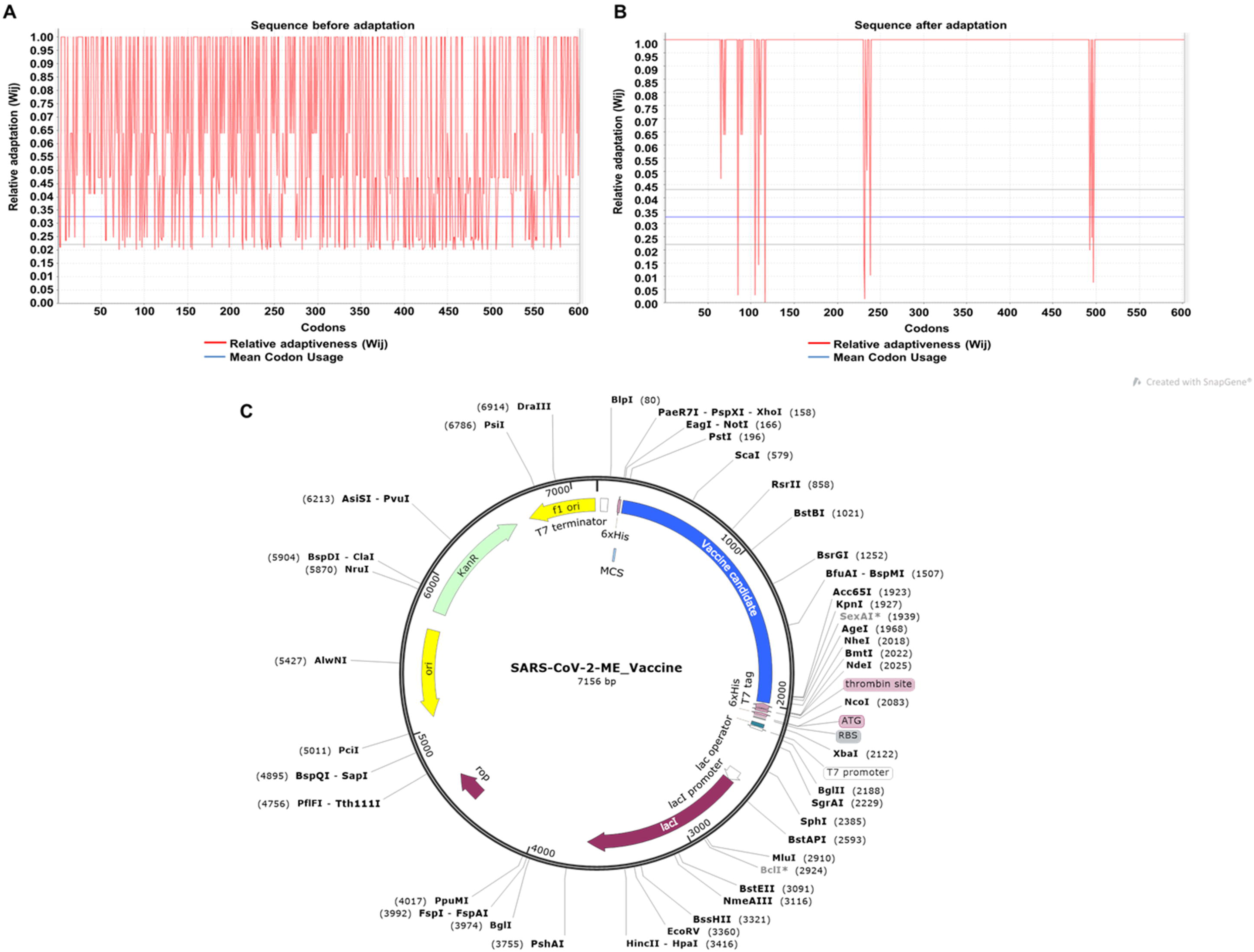
Codon optimization and *In silico* cloning of the cDNA of SARS-CoV-2 multi-epitope vaccine construct. Representation of CAI score of the cDNA construct of SARS-CoV-2 multi-epitope vaccine before adaptation (A) and after adaptation (B). (C) Pictorial representation of SARS-CoV-2-multi-epitope vaccine construct plasmid that can be used for the expression and purification of the vaccine product inside *Escherichia coli*.

### 12. Expression of Multi-epitope vaccine construct in *Escherichia coli* by *In silico* cloning

*Escherichia coli K12* strain was selected as a host for cloning purpose as the expression and purification of multi-epitope vaccines are easier in this bacterium. pET28a(+) expression vector was cleaved using *BamHI* and *HindIII* restriction enzyme and the cDNA was inserted near the ribosome binding site using Snapgene. 6X histidine tag was added at the 3’ end for the isolation and purification of the vaccine construct (Figure 9C).

## DISCUSSION

SARS-CoV-2 that causes COVID-19 has recently emerged as one of the deadliest pathogens severely affecting humans worldwide. The SARS-CoV-2 is highly contagious virus with a high mortality rate especially in immunocompromised and elderly persons. Vaccines are the utmost need of the time to succumb the rising infections due to COVID-19 and represent the best way to combat infectious diseases in the community. Conventional vaccine development methods are time-consuming, laborious and expensive and in this regard, the in-silico immuno-informatics approach have proved to a boon[59]. Moreover, the availability of large number of efficient tools to predict the immuno-determinants, and large database of information facilitate and accelerate this whole process of vaccine designing[60]. Multi-epitope vaccines are designed by using the most antigenic conserved part of the pathogen’s proteins that makes them more effective and can help in combating the high mutation rate in RNA viruses. These epitope-based vaccines have added advantages of being safe, stable, highly specific, cost-curtailing, easy to produce in bulk, and provides option to manipulate the epitopes for designing of a better vaccine candidate[61]. This method of vaccine development has proved to be promising enough for different microbial diseases including both bacterial and viral[25,62–67].

Herein, we applied reverse vaccinology approach for designing of a multi-epitope vaccine that can efficiently elicit humoral and cellular mediated immune response against SARS-CoV-2 viral infection. Coronaviruses are the viruses containing one of the largest RNA genomes encoding several accessory proteins, non-structural proteins and the structural proteins. Generation of humoral and cellular mediated memory immunity leads to fast and long lasting immune response in case of viral infection. In our vaccine development process against the SARS-CoV-2, the antigenic viral proteins were screened for the most immunogenic and antigenic B-cell and T-cell epitopes that could efficiently evoke the immune response. For the generation of most efficacies vaccine, successful recognition of epitopes by various HLA alleles is a preliminary requisite. Therefore, 14 HLA I and 20 HLA II alleles were selected that covers the maximum population in the severely affected countries throughout the world. The screened epitopes were further checked for allergenicity and toxicity inside the host. This screened epitopes covered fifteen various antigenic proteins of SARS-CoV-2 including various non-structural [nsp2, nsp4, nsp-6, Guanine-N7 methyltransferase (ExoN), etc.], structural (E, M, N and S) and accessory proteins (ORF3a, ORF7a and ORF8). Inclusion of all these proteins makes it a potent and effective vaccine candidate against the SARS-CoV-2 virus. The protein–peptide docking of the HL Aalleles and selected epitopes showed the efficient binding in the epitope binding cleft of HLA structure. The population coverage analysis of the vaccine based on the presence of epitopes for the HLA alleles showed the broad range of the vaccine construct covering >99% of world population. Adjuvants are the small peptides that when added along with the antigenic vaccine enhances the immunogenic response. Considering this fact, four different adjuvants were added to the multi-epitope vaccine construct that enhanced the innate and adaptive immune response and helped in better transportation through the intestinal membrane barrier. Vaccine construct was observed to be antigenic, non-allergenic and non-toxic making it a potent vaccine candidate against SARS-CoV-2. The physiochemical analysis of the construct revealed the stable nature of the vaccine product and is at par with the standards required for the formulation of an effective vaccine. The solubility analysis of the multi-epitope vaccine construct revealed ~70 % of solubility in an overexpressed state which is in admissible limits and is required for purification and other functional analysis.

Secondary structure prediction revealed the predominance of alpha helix while the tertiary structure obtained from threading provides the spatial arrangement of amino-acid in space. The obtained modelled structure was further refined that increased its overall quality. To check the interaction of the vaccine construct with the immune receptors, protein-protein docking was performed. Various immune receptors like Toll-like receptors (TLRs) act as sensor proteins that help in recognizing and differentiating self and non-self molecules and cells. They help in recognizing molecular patterns on the surface of foreign particles like viruses, bacteria and fungi and elicit immune response against these infectious agents. TLR3 acts as a nucleic acid sensor and helps in recognizing RNA and DNA viruses inside the human body and establishes a protective role against this infectious agents[68]. In SARS-CoV infection, TLR3 signaling contributes to the innate immune response against the viral infection[69]. Thus agonist of TLR3, β-defensin that also acts as an adjuvant was added to the vaccine construct. Also, the molecular interaction analysis of the vaccine construct with the TLR3 resulted into a stable complex formation that was energetically favorable. RSMD analysis of the vaccine construct and the vaccine construct-TLR3 complex depicted the stable nature of both the systems in the real environment.

Finally, codon optimization was performed for the maximal expression of multi-epitope vaccine product in *Escherichia coli K12*. Overall, herein, an integrated next generation vaccinology approach was used to design a potential antigenic multi-epitope subunit vaccine which may combat the COVID-19 with high specificity and efficacy.

## CONCLUSION

The emergence of novel strain of Corona virus, SARS-CoV-2 has become one of the most detrimental pathogen world-wide. With its high contagious nature and no effective vaccines and drugs, it has become a pandemic throughout. Herein, we explored next generation vaccinology approach for designing multi-epitope vaccine construct. The proteins of SARS-CoV-2 were screened for highly conserved, antigenic, non-allergen and non-toxic HTL, CTL and BCL epitopes. Molecular interaction analysis revealed the high propensity of interaction of the predicted epitopes with HLA alleles. Further *in-silico* analysis revealed the formation of generation of both humoral and cellular mediated immune response. Various physiochemical analyses revealed the formation of stable and efficacious vaccine candidate with a population coverage of >99 % world-wide. Finally, *in-silico* analysis was performed so as to easily express and purify the multi-epitope vaccine construct in a bacterial host. Thus, the present study scrutinizes the antigenic proteins of SARS-CoV-2 for designing and highly efficacious protective vaccine candidate that can elicit both humoral and cellular mediated memory response and provides a starting platform for *in-vitro* and *in-vivo* analysis for the development of protective and prophylactic solution against COVID-19.

## Supplementary Information

Supplementary Figure S1-S5, Supplementary Table S1-S4 and Supplementary Data S1

## Author Contributions

Data conceptualization and methodology was performed by AK. *In silcio* prediction were performed by NJ, US and PM. Analysis was performed by NJ. NJ and PM collectively wrote the manuscript. A.K. did the review and editing.

## Acknowledgments

The authors acknowledge Computer server facility provided by Dr. Parimal Kar at IIT, Indore. NJ acknowledges the Council for Scientific and Industrial Research, Govt. of India, New Delhi, while US and PM acknowledges Ministry of Human Resource and Development, Govt. of India, New Delhi and for respective Ph.D. research fellowships.

