## Supplementary Figure, Supplementary Table for "Scrutinizing the SARS-CoV-2 protein information for the designing an effective vaccine encompassing both the T-cell and B-cell epitopes"

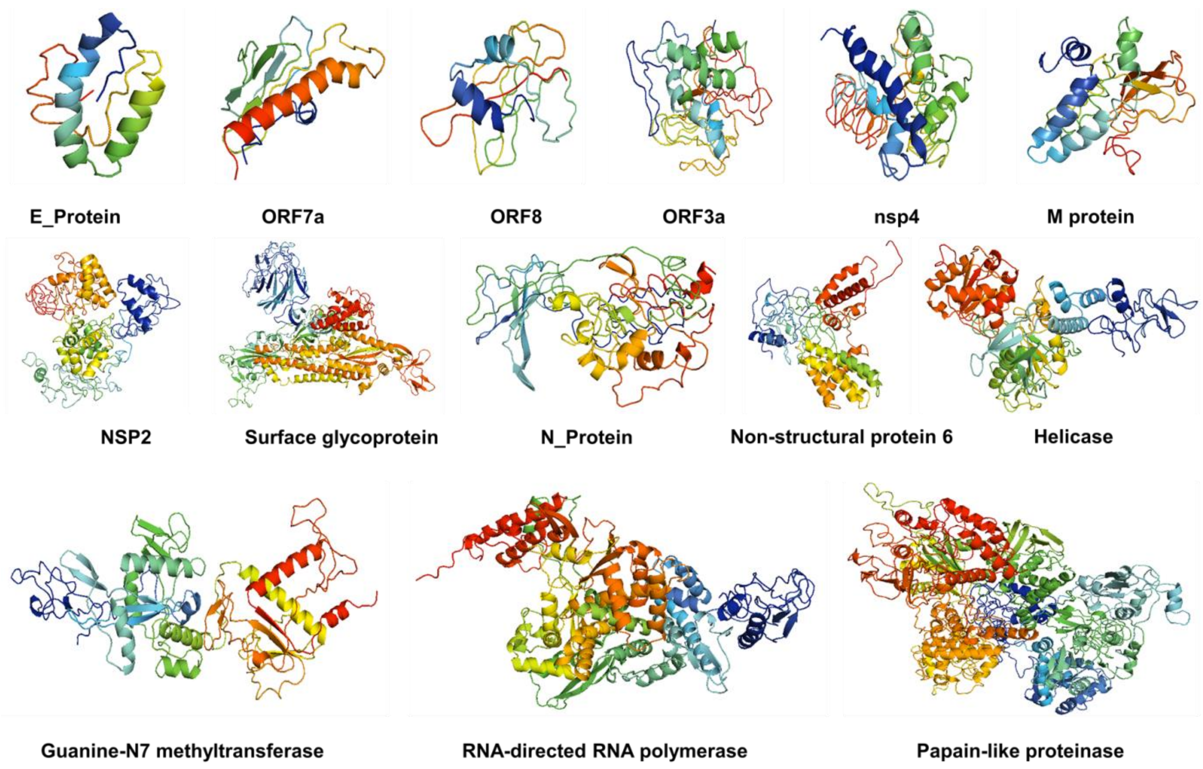


Supplementary Figure S1. 3 dimensional structures in ribbon illustration of the modelled SARS-CoV-2 proteins as predicted by I-TASSER. The Figures are generated using Pymol visualization tool.

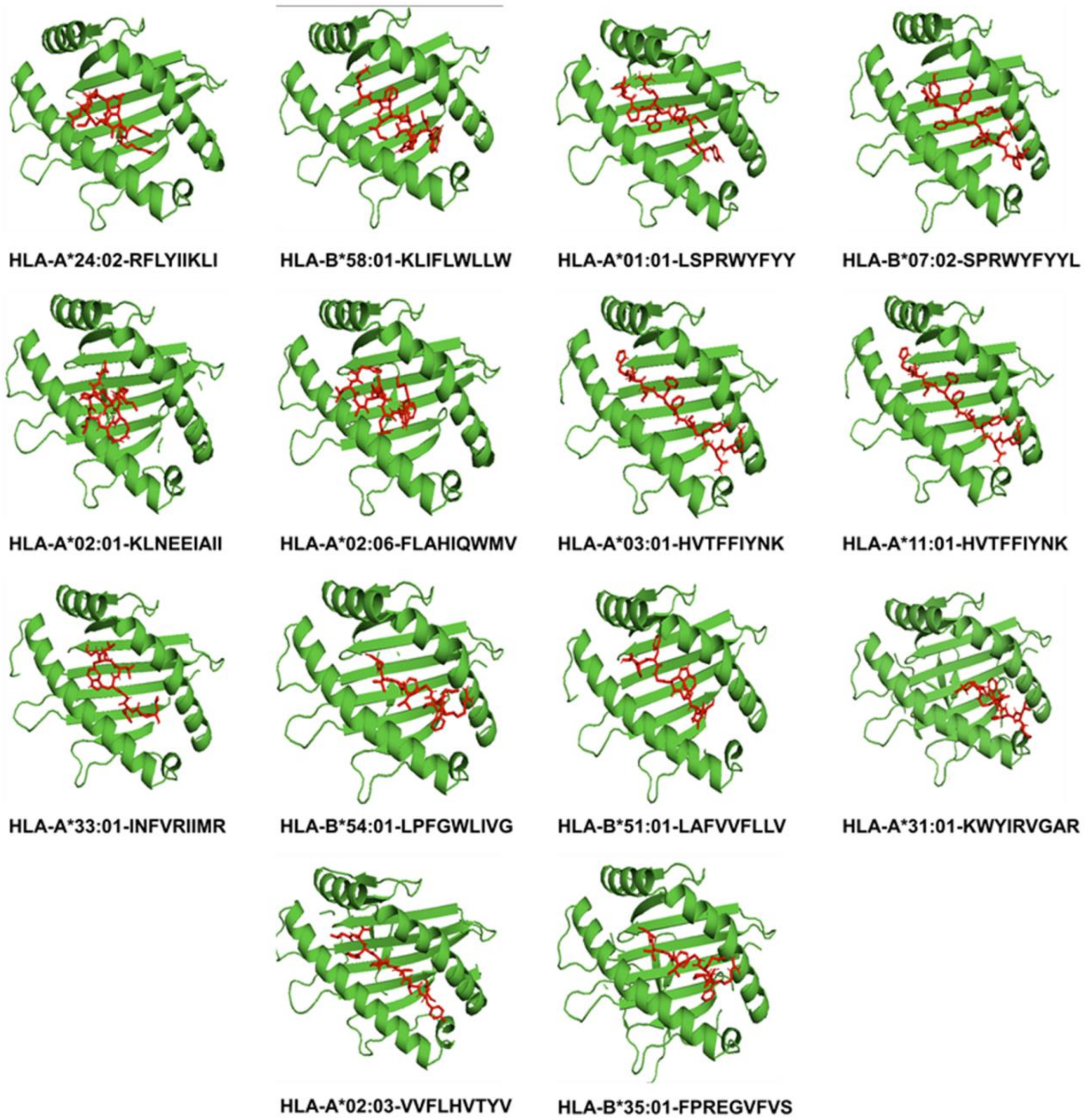


**Supplementary Figure S2: The molecular interaction of the screened epitopes with the epitope binding groove of HLA molecules.**

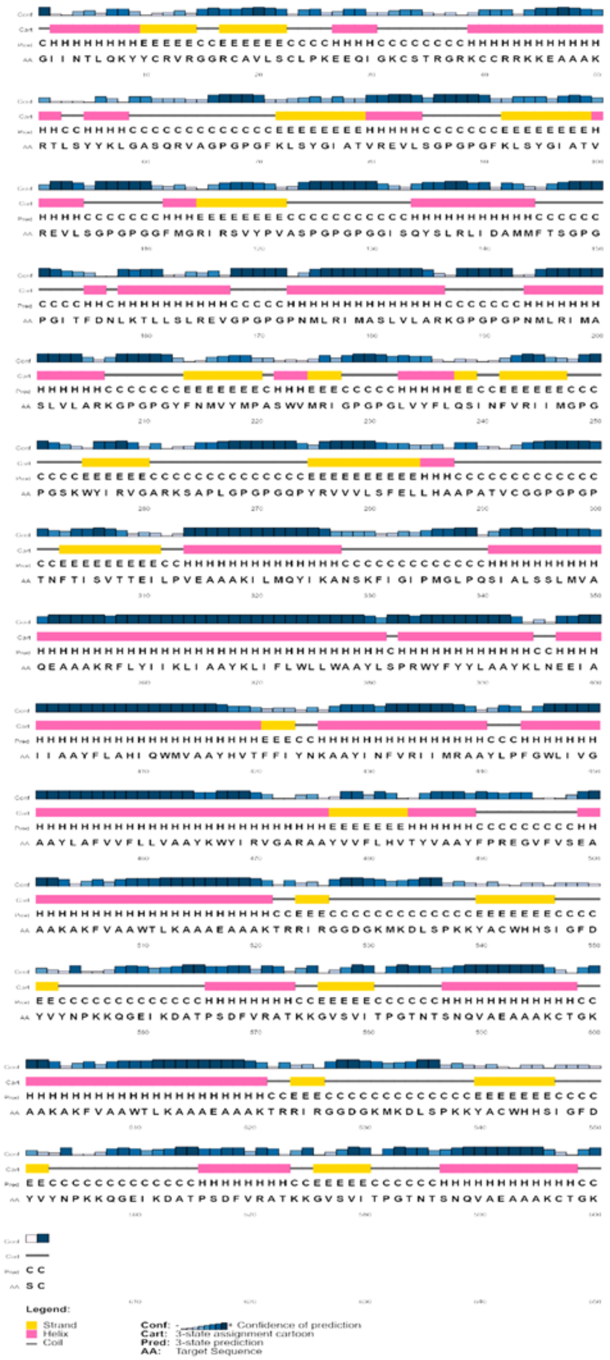

**Supplementary Figure S3. Secondary Structure prediction result of the SARS-CoV-2 multi-epitope vaccine as predicted by PSI-PRED server.**

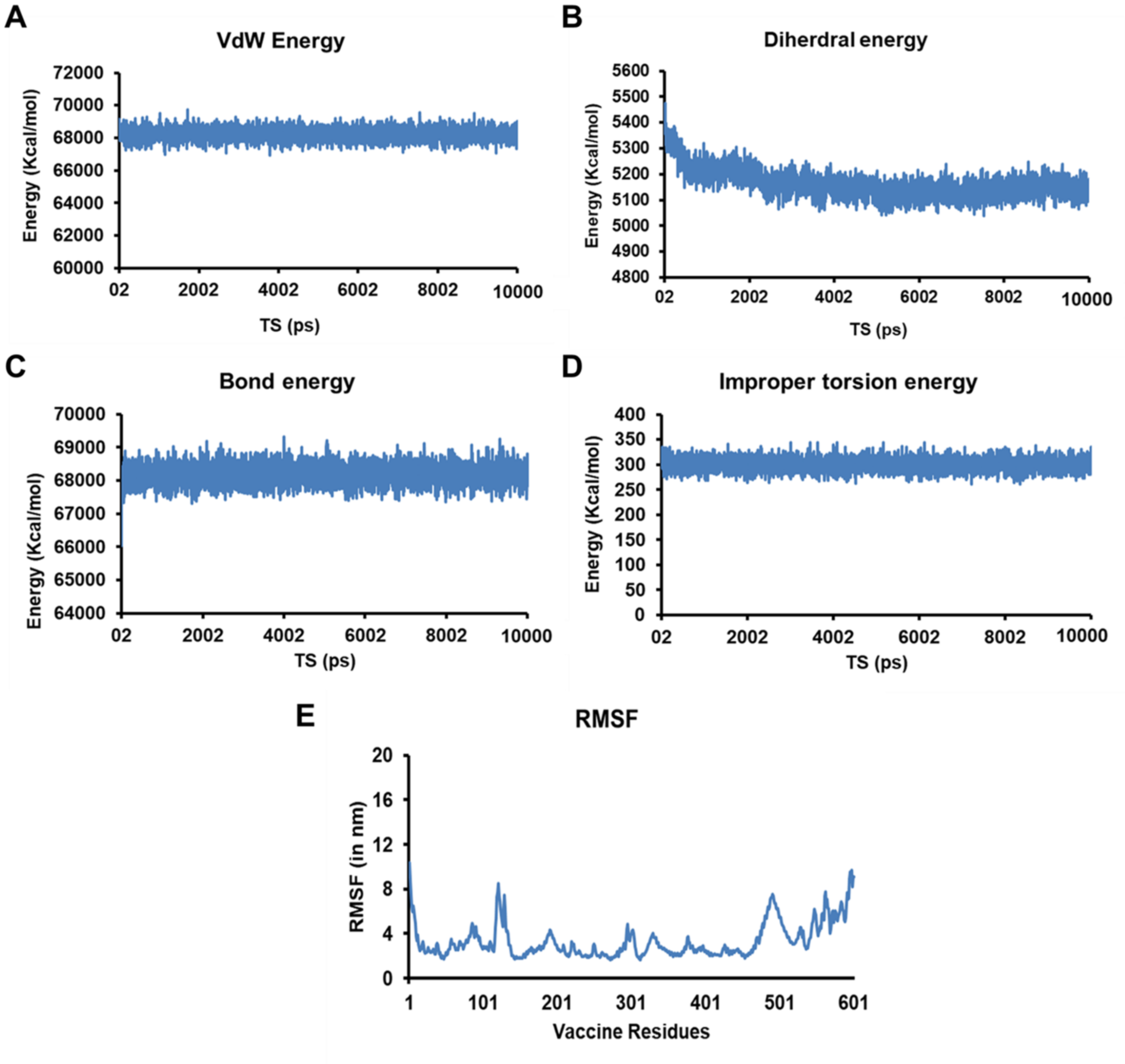


**Supplementary Figure S4. (A-D) Energy plots depicting the vanderwaal's energy (A), dihedral angle energy (B), improper dihedral energy (C) and bond energy (D) of the SARS-CoV-2 multi-epitope vaccine during the 10 ns molecular dynamic simulation analysis. (E) Root mean square fluctuation as observed in the residues of the vaccine construct in 10ns duration.**

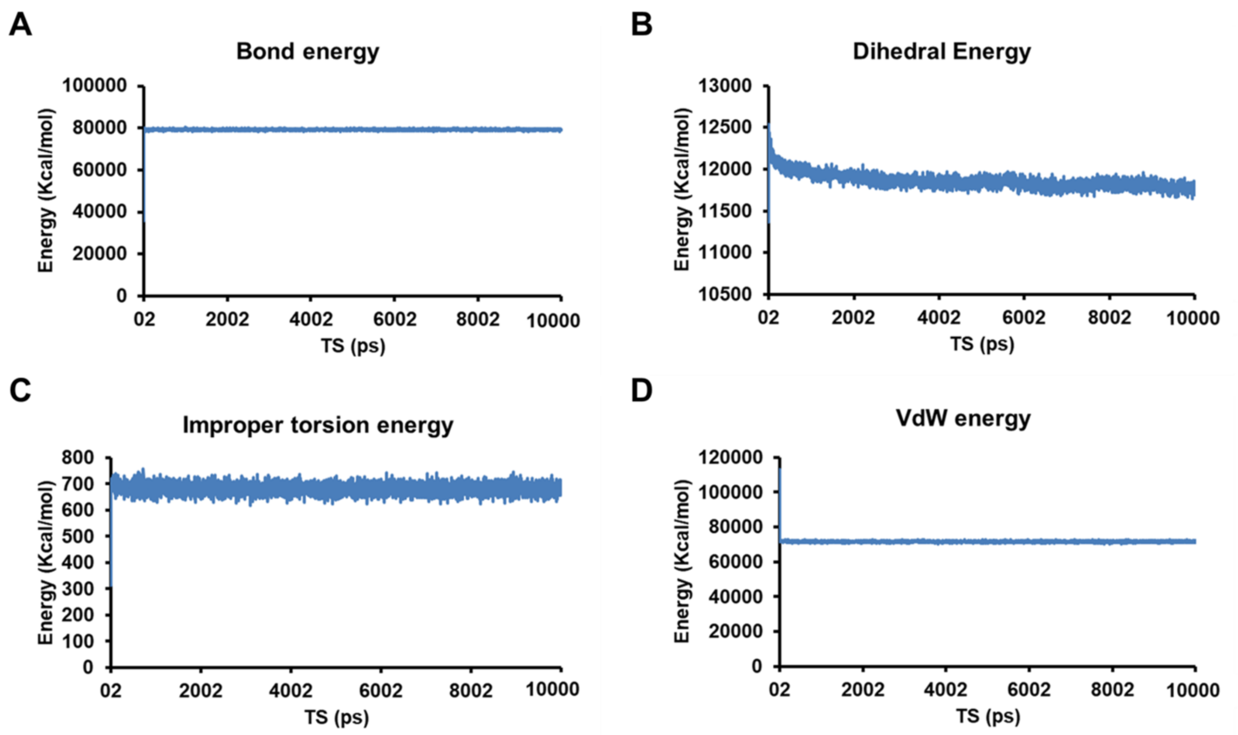


**Supplementary Figure S5. (A-D) Energy plots depicting the vanderwaal's energy (A), dihedral angle energy (B), improper dihedral energy (C) and bond energy (D) of the SARS-CoV-2 multi-epitope vaccine and TLR3 complex during the 10 ns molecular dynamic simulation analysis.**

**Supplementary Table S1. List of Helper T-cell epitopes of SARS-CoV-2 antigenic proteins selected for the multi-epitope vaccine construct with the percentile rank, antigenicity score and allergenicity prediction.**

| SARS-CoV-2 proteins | Position | HLA alleles | Method | Epitope Sequence | percentile_rank | adjusted_rank | Antigenicity Score | Allergenicity (AlgPred) |
|---|---|---|---|---|---|---|---|---|
| M glycoprotein | 174 | HLA-DRB1*09:01 | Consensus (comb.lib./smm/nn) | RTLSYYKLGASQRVA | 0.06 | 0.06 | 0.5644 | NON ALLERGEN |
| ORF1ab [Helicase (Hel)] | 5469 | HLA-DQA1*04:01/DQB1*04:01 | NetMHCIIpan | FKLSYGIATVREVLS | 0.89 | 0.89 | 0.7783 | NON ALLERGEN |
| ORF1ab [Helicase (Hel)] | 5469 | HLA-DQA1*04:02/DQB1*04:02 | NetMHCIIpan | FKLSYGIATVREVLS | 0.89 | 0.89 | 0.7783 | NON ALLERGEN |
| ORF1ab (Non-structural protein 2_nsp2) | 295 | HLA-DRB1*08:02 | Consensus (smm/nn/sturniolo) | GFMGRIRSVYPVASP | 0.64 | 0.64 | 0.7131 | NON ALLERGEN |
| ORF1ab(Non-structural protein 2_nsp2) | 572 | **HLA-DQA1*01:01/DQB1*02:01** | NetMHCIIpan | GISQYSLRLIDAMMF | 0.66 | 0.66 | 0.7195 | NON ALLERGEN |
| ORF1ab (Non-structural protein 2_nsp2) | 574 | HLA-DQA1*01:02/DQB1*02:02 | NetMHCIIpan | SQYSLRLIDAMMFTS | 0.33 | 0.33 | 0.7276 | NON ALLERGEN |
| ORF1ab Non-structural protein 6 (nsp6) | 3649 | HLA-DQA1*05:01/DQB1*05:01 | NetMHCIIpan | YFNMVYMPASWVMRI | 0.19 | 0.19 | 0.7244 | NON ALLERGEN |
| ORF1ab RNA-directed RNA polymerase (RdRp) | 5019 | HLA-DPA1*03:01/DPB1*03:01 | NetMHCIIpan | PNMLRIMASLVLARK | 0.06 | 0.06 | 0.4128 | NON ALLERGEN |
| ORF1ab RNA-directed RNA polymerase (RdRp) | 5019 | HLA-DRB1*15:01 | Consensus (smm/nn/sturniolo) | PNMLRIMASLVLARK | 0.01 | 0.01 | 0.4128 | NON ALLERGEN |
| ORF1ab Papain-like proteinase | 1551 | HLA-DRB1*04:01 | Consensus (smm/nn/sturniolo) | ITFDNLKTLLSLREV | 0.33 | 0.33 | 0.8234 | NON ALLERGEN |
| ORF3a protein | 111 | HLA-DPA1*02:01/DPB1*02:01 | NetMHCIIpan | LVYFLQSINFVRIIM | 0.09 | 0.09 | 0.5186 | NON ALLERGEN |
| ORF8 protein | 43 | HLA-DRB1*11:01 | Consensus (smm/nn/sturniolo) | SKWYIRVGARKSAPL | 0.72 | 0.72 | 0.8829 | NON ALLERGEN |
| S protein | 506 | HLA-DPA1*01:03/DPB1*01:01 | NetMHCIIpan | **QPYRVVVLSFELLHA** | 0.29 | 0.29 | 0.9109 | NON ALLERGEN |
| S protein | 512 | HLA-DRB1*01:01 | Consensus (comb.lib./smm/nn) | **VLSFELLHAPATVCG** | 0.03 | 0.03 | 0.4784 | NON ALLERGEN |
| S protein | 715 | HLA-DRB1*07:01 | Consensus (comb.lib./smm/nn) | PTNFTISVTTEILPV | 0.51 | 0.51 | 1.1349 | NON ALLERGEN |

**Supplementary Table S2. List of Cytotoxic T-cell epitopes of SARS-CoV-2 antigenic proteins selected for the multi-epitope vaccine construct with the percentile rank, binding level, immunogenicity score, antigenicity score and allergenicity prediction.**

| SARS-CoV-2 Protein | Start genomic point | HLA Allele | Epitope | Score | Aff(nM) | %Rank | Binding Level | Antigencity score | Immunogenecity Score | Allergenicity |
|---|---|---|---|---|---|---|---|---|---|---|
| M Protein | 44 | HLA-A*24:02 | RFLYIIKLI | 0.496296 | 232.8 | 0.4035 | SB | 0.4257 | 0.24108 | PROBABLE NON-ALLERGEN The nearest protein is: UniProtKB accession number O00626 defined as non-allergen |
| M Protein | 50 | HLA-B*58:01 | KLIFLWLLW | 0.588682 | 85.7 | 0.4281 | SB | 0.4968 | 0.34287 | PROBABLE NON-ALLERGEN The nearest protein is: UniProtKB accession number P60140 defined as non-allergen |
| N Protein | 104 | HLA-A*01:01 | LSPRWYFYY | 0.59866 | 76.9 | 0.073 | SB | 1.2832 | 0.36094 | PROBABLE NON-ALLERGEN The nearest protein is: UniProtKB accession number Q0VD83 defined as non-allergen |
| N Protein | 105 | HLA-B*07:02 | SPRWYFYYL | 0.748063 | 15.3 | 0.0496 | SB | 0.734 | 0.34101 | PROBABLE NON-ALLERGEN The nearest protein is: UniProtKB accession number Q7XAQ6 defined as non-allergen |
| ORF1ab (Non-structural protein 2_nsp2) | 468 | HLA-A*02:01 | KLNEEIAI | 0.673256 | 34.3 | 0.4552 | SB | 0.6394 | 0.43221 | PROBABLE NON-ALLERGEN The nearest protein is: UniProtKB accession number A2Z5D8defined as non-allergen |
| ORF1ab (Non-structural protein 4 (nsp4) | 3122 | HLA-A*02:06 | FLAHIQWMV | 0.914969 | 2.5 | 0.0074 | SB | 0.8064 | 0.38891 | PROBABLE NON-ALLERGEN The nearest protein is: UniProtKB accession number P31358 defined as non-allergen |
| ORF3a Protein | 227 | HLA-A*03:01 | HVTFFIYNK | 0.544282 | 138.5 | 0.4821 | SB | 0.9862 | 0.36278 | PROBABLE NON-ALLERGEN The nearest protein is: UniProtKB accession number O43852 defined as non-allergen |
| ORF3a Protein | 227 | HLA-A*11:01 | HVTFFIYNK | 0.759484 | 13.5 | 0.0517 | SB | 0.9862 | 0.36278 | PROBABLE NON-ALLERGEN The nearest protein is: UniProtKB accession number O43852 defined as non-allergen |
| ORF3a Protein | 118 | HLA-A*33:01 | INFVRIIMR | 0.538132 | 148 | 0.3814 | SB | 0.7646 | 0.26494 | PROBABLE NON-ALLERGEN The nearest protein is: UniProtKB accession number O15511 defined as non-allergen |
| ORF3a Protein | 41 | HLA-B*54:01 | LPFGWLIVG | 0.63608 | 51.3 | 0.0762 | SB | 1.981 | 0.4138 | PROBABLE NON-ALLERGEN The nearest protein is: UniProtKB accession |

| | | | | | | | | | | |
|---|---|---|---|---|---|---|---|---|---|---|
| | | | | | | | | | | number Q156A1 defined as non-allergen |
| ORF4 (E protein) | 21 | HLA-B*51:01 | LAFVVFL LV | 0.305577 | 1832.6 | 0.3161 | SB | 0.7976 | 0.2141 | PROBABLE NON-ALLERGEN The nearest protein is: UniProtKB accession number P60140 defined as non-allergen |
| ORF8 Protein | 44 | HLA-A*31:01 | KWYIRV GAR | 0.72322 | 20 | 0.1425 | SB | 1.5772 | 0.27344 | PROBABLE NON-ALLERGEN The nearest protein is: UniProtKB accession number Q96PC3 defined as non-allergen |
| S Protein | 1060 | HLA-A*02:03 | VVFLHVT YV | 0.793601 | 9.3 | 0.1844 | SB | 1.5122 | 0.1278 | PROBABLE NON-ALLERGEN The nearest protein is: UniProtKB accession number Q6UWT4 defined as non-allergen |
| S Protein | 1089 | HLA-B*35:01 | FPREGVF VS | 0.56001 | 116.8 | 0.3526 | SB | 0.5509 | 0.31233 | PROBABLE NON-ALLERGEN The nearest protein is: UniProtKB accession number P07711 defined as non-allergen |

**Supplementary Table S3. List of B cell Lymphocytes epitopes of SARS-CoV-2 antigenic proteins selected for the multi-epitope vaccine construct with the ABCPred prediction score, antigenicity score and allergenicity prediction.**

| SARS-CoV-2 proteins | B cell epitope Sequence | Start position | Score | Antigenicity Score | AllerTop V2 |
|---|---|---|---|---|---|
| Nucleocapsid | TRRIRGGDGKMKDLSP | 91 | 0.94 | 1.1467 | Non-allergen |
| ORF1ab Guanine-N7 methyltransferase (ExoN) | YACWHHSIGFDYVYNP | 6149 | 0.96 | 0.9219 | Non-allergen |
| ORF3a | QGEIKDATPSDFVRAT | 17 | 0.94 | 0.9161 | Non-allergen |
| Surface glycoprotein | GVSVITPGTNTSNQVA | 594 | 0.95 | 0.4651 | Non-allergen |

**Supplementary Table S4. The Lowest energy score obtained for the 30 conformers of TLR3 – Vaccine construct complex as predicted by ClusPro results.**

| Cluster | Members | Representative | Weighted Score |
|---|---|---|---|
| 0 | 35 | Center | -1227.0 |
| | | Lowest Energy | -1275.4 |
| 1 | 30 | Center | -1106.4 |
| | | Lowest Energy | -1171.4 |
| 2 | 29 | Center | -1198.2 |
| | | Lowest Energy | -1198.2 |
| 3 | 25 | Center | -1036.6 |
| | | Lowest Energy | -1158.1 |
| 4 | 23 | Center | -1108.9 |
| | | Lowest Energy | -1108.9 |
| 5 | 21 | Center | -999.0 |
| | | Lowest Energy | -1427.4 |
| 6 | 21 | Center | -1243.7 |
| | | Lowest Energy | -1243.7 |
| 7 | 20 | Center | -1030.6 |
| | | Lowest Energy | -1128.3 |
| 8 | 19 | Center | -1101.4 |
| | | Lowest Energy | -1146.2 |
| 9 | 18 | Center | -1252.6 |
| | | Lowest Energy | -1287.4 |
| 10 | 17 | Center | -1202.9 |
| | | Lowest Energy | -1202.9 |
| 11 | 16 | Center | -1017.1 |
| | | Lowest Energy | -1150.9 |
| 12 | 16 | Center | -1218.4 |

| | | | |
|---|---|---|---|
| | | Lowest Energy | -1218.4 |
| 13 | 16 | Center | -1052.3 |
| | | Lowest Energy | -1140.9 |
| 14 | 16 | Center | -1012.1 |
| | | Lowest Energy | -1268.0 |
| 15 | 15 | Center | -1032.3 |
| | | Lowest Energy | -1356.2 |
| 16 | 15 | Center | -1156.2 |
| | | Lowest Energy | -1266.9 |
| 17 | 15 | Center | -1131.0 |
| | | Lowest Energy | -1267.3 |
| 18 | 15 | Center | -1069.0 |
| | | Lowest Energy | -1162.9 |
| 19 | 14 | Center | -1014.0 |
| | | Lowest Energy | -1160.3 |
| 20 | 13 | Center | -1189.0 |
| | | Lowest Energy | -1189.0 |
| 21 | 13 | Center | -1130.0 |
| | | Lowest Energy | -1130.0 |
| 22 | 13 | Center | -1084.9 |
| | | Lowest Energy | -1084.9 |
| 23 | 13 | Center | -1055.6 |
| | | Lowest Energy | -1070.5 |
| 24 | 12 | Center | -1084.9 |
| | | Lowest Energy | -1084.9 |
| 25 | 11 | Center | -1036.2 |
| | | Lowest Energy | -1123.7 |
| 26 | 11 | Center | -1205.6 |
| | | Lowest Energy | -1205.6 |
| 27 | 11 | Center | -1119.2 |
| | | Lowest Energy | -1119.2 |
| 28 | 11 | Center | -1098.0 |
| | | Lowest Energy | -1206.7 |
| 29 | 11 | Center | -1019.6 |
| | | Lowest Energy | -1077.6 |

**Supplementary Data S1. Codon optimized and adapted cDNA sequence of the SARD-CoV multi-epitope vaccine construct.**

GGTATCATCAACACCCTGCAGAAATACTACTGCCGTGTTCGTGGTGGTCG 50
TTGCGCTGTTCTGTCTTGCCTGCCGAAAGAAGAACAGATCGGTAAATGCT 100
CTACCCGTGGTCGTAAATGCTGCCGTCGTAAAAAAGAAGCTGCTGCTAAA 150
CGTACCCTGTCTTACTACAAACTGGGTGCTTCTCAGCGTGTTGCGGGTCC 200
GGGCCCGGGCTTCAAACTGTCTTACGGTATCGCTACCGTTCGTGAAGTTC 250
TGTCGGGTCCGGGCCCGGGCTTCAAACTGTCTTACGGTATCGCTACCGTT 300
CGTGAAGTTCTGTCGGGTCCGGGTCCAGGTGGCTTCATGGGTCGTATACG 350
TTCTGTTTACCCGGTTGCTTCTCCGGGTCCGGGTCCGGGTGGTATCTCTC 400
AGTACTCTCTGCGTCTGATCGACGCTATGATGTTCACCTCTGGTCCGGGT 450
CCGGGTATCACCTTCGACAACCTGAAAACCCTGCTGTCTCTGCGTGAAGT 500
TGGTCCGGGTCCGGGTCCGAACATGCTGCGTATCATGGCTTCTCTGGTTC 550
TGGCTCGTAAAGGTCCGGGTCCGGGTCCGAACATGCTGCGTATCATGGCT 600
TCTCTGGTTCTGGCTCGTAAAGGTCCGGGTCCGGGTTACTTCAACATGGT 650
TTACATGCCGGCTTCTTGGGTTATGCGTATCGGTCCGGGTCCAGGGCTGG 700
TATACTTCCTGCAATCTATCAACTTCGTTCGTATCATCATGGGTCCGGGT 750
CCGGGTTCTAAATGGTACATCCGTGTTGGTGCTCGTAAATCTGCTCCGCT 800
GGGTCCGGGTCCGGGTCAGCCGTACCGTGTTGTTGTTCTGTCTTTCGAAC 850
TGCTGCACGCTGCTCCGGCTACCGTTTGCGGTGGTCCGGGTCCGGGTCCG 900
ACCAACTTCACCATCTCTGTTACCACCGAAATCCTGCCGGTTGAAGCTGC 950
TGCTAAAATCCTGATGCAGTACATCAAAGCTAACTCTAAATTCATCGGTA 1000
TCCCGATGGGTCTGCCGCAGTCTATCGCTCTGTCTTCTCTGATGGTTGCT 1050
CAGGAAGCTGCTGCTAAACGTTTCCTGTACATCATCAAACTGATCGCTGC 1100
TTACAAACTGATCTTCCTGTGGCTGCTGTGGGCTGCTTACCTGTCTCCGC 1150
GTTGGTACTTCTACTACCTGGCTGCTTACAAACTGAACGAAGAAATCGCT 1200
ATCATCGCTGCTTACTTCCTGGCTCACATCCAGTGGATGGTTGCTGCTTA 1250
CCACGTTACCTTCTTCATCTACAACAAAGCTGCTTACATCAACTTCGTTC 1300
GTATCATCATGCGTGCTGCTTACCTGCCGTTCGGTTGGCTGATCGTTGGT 1350
GCTGCTTACCTGGCTTTCGTTGTTTTCCTGCTGGTTGCTGCTTACAAATG 1400
GTACATCCGTGTTGGTGCTCGTGCTGCTTACGTTGTTTTCCTGCACGTTA 1450
CCTACGTTGCTGCTTACTTCCCGCGTGAGGGTGTGTTCGTCTCTGAAGCT 1500
GCTGCTAAAGCTAAATTCGTTGCTGCTTGGACCCTGAAAGCTGCTGCTGA 1550
AGCTGCTGCTAAAACCCGTCGTATCCGTGGTGGTGACGGTAAAATGAAAG 1600
ACCTGTCTCCGAAAAAATACGCTTGCTGGCACCACTCTATCGGTTTCGAC 1650
TACGTTTACAACCCGAAAAAACAGGGTGAAATCAAAGACGCTACCCCGTC 1700
TGACTTCGTTCGTGCTACCAAAAAAGGTGTTTCTGTTATCACCCCGGGTA 1750
CCAACACCTCTAACCAGGTTGCTGAAGCTGCTGCTAAATGCACCGGTAAA 1800
TCTTGC
